# Genetic and structural basis for recognition of SARS-CoV-2 spike protein by a two-antibody cocktail

**DOI:** 10.1101/2021.01.27.428529

**Authors:** Jinhui Dong, Seth J. Zost, Allison J. Greaney, Tyler N. Starr, Adam S. Dingens, Elaine C. Chen, Rita E. Chen, James Brett Case, Rachel E. Sutton, Pavlo Gilchuk, Jessica Rodriguez, Erica Armstrong, Christopher Gainza, Rachel S. Nargi, Elad Binshtein, Xuping Xie, Xianwen Zhang, Pei-Yong Shi, James Logue, Stuart Weston, Marisa E. McGrath, Matthew B. Frieman, Tyler Brady, Kevin Tuffy, Helen Bright, Yueh-Ming Loo, Patrick McTamney, Mark Esser, Robert H. Carnahan, Michael S. Diamond, Jesse D. Bloom, James E. Crowe

**Affiliations:** Vanderbilt Vaccine Center, Vanderbilt University Medical Center, Nashville, TN 37232, USA; Basic Sciences Division, Fred Hutchinson Cancer Research Center, Seattle, WA 98109, USA; Department of Genome Sciences & Medical Scientist Training Program, University of Washington, Seattle, WA 98195, USA; Department of Pathology, Microbiology, and Immunology, Vanderbilt University Medical Center, Nashville, TN, 37232, USA; Department of Pathology and Immunology, Washington University School of Medicine, Saint Louis, MO, 63110, USA; Department of Medicine, Washington University School of Medicine, Saint Louis, MO, 63110, USA; Department of Biochemistry & Molecular Biology, The University of Texas Medical Branch at Galveston, Galveston, TX, 77555, USA; Department of Microbiology and Immunology, The University of Maryland, College Park, MD, 20742, USA; Microbial Sciences, AstraZeneca, One MedImmune Way, Gaithersburg, MD 20878, USA; Department of Pediatrics, Vanderbilt University Medical Center, Nashville, TN, 37232, USA; Department of Molecular Microbiology, Washington University School of Medicine, Saint Louis, MO, 63110, USA; Andrew M. and Jane M. Bursky Center for Human Immunology and Immunotherapy Programs, Washington University School of Medicine, Saint Louis, MO, 63110, USA; Howard Hughes Medical Institute, Seattle, WA, 98109, USA

**Author notes:** Correspondence to: James E. Crowe, Jr., M.D., Contact information: James E. Crowe, Jr., M.D. [LEAD CONTACT], Departments of Pediatrics, Pathology, Microbiology, and Immunology, and the Vanderbilt Vaccine Center, Mail: Vanderbilt Vaccine Center, 11475 Medical Research Building IV 2213 Garland Avenue Nashville, TN 37232-0417, USA. These authors contributed equally (Jinhui Dong, Seth J. Zost).

**Keywords:** Coronavirus, SARS-CoV-2, SARS-CoV, COVID-19, Antibodies, Monoclonal, Human, Adaptive Immunity.

## Abstract

The SARS-CoV-2 pandemic has led to an urgent need to understand the molecular basis for immune recognition of SARS-CoV-2 spike (S) glycoprotein antigenic sites. To define the genetic and structural basis for SARS-CoV-2 neutralization, we determined the structures of two human monoclonal antibodies COV2-2196 and COV2-2130^1^, which form the basis of the investigational antibody cocktail AZD7442, in complex with the receptor binding domain (RBD) of SARS-CoV-2. COV2-2196 forms an “aromatic cage” at the heavy/light chain interface using germline-encoded residues in complementarity determining regions (CDRs) 2 and 3 of the heavy chain and CDRs 1 and 3 of the light chain. These structural features explain why highly similar antibodies (public clonotypes) have been isolated from multiple individuals^1–4^. The structure of COV2-2130 reveals that an unusually long LCDR1 and HCDR3 make interactions with the opposite face of the RBD from that of COV2-2196. Using deep mutational scanning and neutralization escape selection experiments, we comprehensively mapped the critical residues of both antibodies and identified positions of concern for possible viral escape. Nonetheless, both COV2-2196 and COV2-2130 showed strong neutralizing activity against SARS-CoV-2 strain with recent variations of concern including E484K, N501Y, and D614G substitutions. These studies reveal germline-encoded antibody features enabling recognition of the RBD and demonstrate the activity of a cocktail like AZD7442 in preventing escape from emerging variant viruses.

---

The current coronavirus disease 2019 (COVID-19) pandemic is caused by SARS-CoV-2, a clade B betacoronavirus (*Sarbecovirus* subgenus) with 96.2% or 79.6% genome sequence identity to the bat coronavirus RaTG13 or SARS-CoV respectively^5, 6^. The S glycoprotein mediates viral attachment via binding to the host receptor angiotensin converting enzyme 2 (ACE2) and possibly other host factors, and subsequent entry into cells after priming by the host transmembrane protease serine 2 (TMPRSS2)^7–9^. The trimeric S protein consists of two subunits, designated S1 and S2. The S1 subunit binds to ACE2 with its receptor binding domain (RBD), while the central trimeric S2 subunits function as a fusion apparatus after S protein sheds the S1 subunits^10^. The human humoral immune response to SARS-CoV-2 has been well documented^11–, 13^, and numerous groups have isolated monoclonal antibodies (mAbs) that react to SARS-CoV-2 S protein from the B cells of patients previously infected with the virus. A subset of the human mAbs neutralize virus *in vitro* and protect against disease in animal models^1, 2, 13–21^. Studies of the human B cell response to the virus have been focused mostly on S protein so far, due to its critical functions in attachment and entry into host cells^1, 2, 13–21^. For these S-protein-targeting antibodies, the RBD of S protein is the dominant target of human neutralizing antibody responses^1, 2, 13–21^. This high frequency of molecular recognition may be related to the accessibility of the RBD to B cell receptors, stemming from a low number of obscuring glycosylation sites (only 2 sites on the RBD versus 8 or 9 sites on the N-terminal domain [NTD] or S2 subunit, respectively)^13^. The RBD also occupies an apical position and exhibits exposure due to the “open-closed” dynamics of the S trimer observed in S protein cryo-EM structures^22–24^. Potently neutralizing mAbs predominantly target the RBD, since this region is directly involved in receptor binding.

In previous studies, we isolated a large panel of SARS-CoV-2 S-protein-reactive human mAbs from the B cells of patients previously infected with the virus. that bind to the SARS-CoV-2 S protein^25^. A subset of these mAbs was shown to bind to recombinant RBD and S protein ectodomain and exhibit neutralization activity against SARS-CoV-2 by blocking S-protein-mediated binding to receptor^1^. Two noncompeting antibodies, designated COV2-2196 and COV2-2130, synergistically neutralized SARS-CoV-2 *in vitro* and protected against SARS-CoV-2 infection in mouse models and a rhesus macaque model when used separately or in combination. Several Phase III clinical trials are ongoing to study AZD7442, which incorporates mAbs that contain the variable regions in this mAb combination, for post-exposure prophylaxis (ClinicalTrials.gov Identifier: NCT04625972), prevention (Identifier: NCT04625725), out-patient treatment (Identifier: NCT04723394 and NCT04518410) and in-patient treatment (NCT04501978) of COVID-19. Thus, it is of importance to define the binding sites of these two antibodies to understand how they interact with the RBD and their ability to neutralize new virus variants.

One of these antibodies (COV2-2196) is a member of a public clonotype, meaning this antibody shares similar variable region genetic features with other antibodies isolated from different individuals. Here, by studying the interaction of COV2-2196 with RBD in detail, we identify the molecular basis for selection of a public clonotype for SARS-CoV-2 that is driven by a complex structural configuration involving both *IGHV1-58-IGHJ3* heavy chain and *IGKV3-20-IGKJ1* light chain recombinations,. The shared structural features of this clonotype contribute to the formation of a paratope comprising residues in both the heavy and light chains, but are independent of the HCDR3 that usually dominates antigen-antibody interactions. Detailed structural studies revealed that the commonly formed antibody paratope contributes an “aromatic cage” formed by five aromatic residues in the paratope surrounding the interface of the heavy and light chains. This cage structure coordinates an aromatic residue on the SARS-CoV-2 S protein, accounting for the high specificity and affinity of these antibodies. Although both the heavy and light chains are required to form this public clonotype (thus defining canonical *IGHV*, *IGHJ*, *IGLV* and *IGLJ* genes in the clonotype), the HCDR3 minimally affects the interaction. Since these *IGHV1-58-IGHJ3* heavy chain and *IGKV3-20-IGKJ1* light chain recombinations are common in the pre-immune B cell repertoire, many individuals likely make such clones during the response to SARS-CoV-2 infection or vaccination. The antigenic site recognized by the complex pre-configured structure of this public clonotype is likely an important component of a protective vaccine for COVID-19 because of the frequency of the B cell clone in the human population and the neutralizing and protective potency of the antibodies encoded by the variable gene segments.

An antibody cocktail including half-life extended versions of COV2-2196 and a non-competing RBD-specific neutralizing antibody, COV2-2130, is being investigated for both prophylaxis and therapy in the trials cited above. To understand the molecular details of the recognition of RBD by COV2-2196 and COV2-2130, we determined the crystal structures of the S protein RBD in complex with COV2-2196 at 2.50 Å (**Fig. 1, Extended Data Table 1**) and in complex with both COV2-2196 and COV2-2130 at 3.00 Å (**Fig. 2, Extended Data Table 1**). The substructure of RBD-COV2-2196 in the RBD-COV2-2196 + COV2-2130 complex is superimposable with that in the structure of the RBD-COV2-2196 complex (**Extended Data Fig. 1**). The buried surface area of the interface between COV2-2196 and the RBD is about 650 Å^2^ in both crystal structures, and that of the interface between COV2-2130 and RBD is about 740 Å^2^. COV2-2196 binds to the receptor-binding ridge of RBD, and COV2-2130 binds to one side of the RBD edge around residue K444 and the saddle region of the receptor binding motif RBM), both partially overlapping the ACE2 binding site (**Fig. 1a-b, 2a-b**). These features explain the competition between the antibodies and ACE2 for RBD binding from our previous study, *e.g*., both COV2-2196 and COV2-2130 neutralize the virus by blocking RBD access to the human receptor ACE2^1^. Aromatic residues from the COV2-2196 heavy and light chains form a hydrophobic pocket that surrounds RBD residue F486 and adjacent residues (G485, N487) (**Fig. 1a, 1d, 1e; Extended Data Fig. 2a-c**). This mode of antibody-antigen interaction is unusual in that the formation of the antibody pocket is caused by wide spatial separation of the HCDR3 and LCDR3. In addition, although the antigenic site recognized by COV2-2196 is not buried at the interface between protomers of S trimer *per se*, COV2-2196 is not able to bind RBD in the “down” conformation due to steric clashes with RBD in an adjacent S protomer. Therefore, COV2-2196 only binds to RBD in the “up” conformation (**Fig. 1c**). Overlays of the substructure of RBD in complex with COV2-2130 (**Fig. 2c)** and the structure of RBD in complex with both COV2-2196 and COV2-2130 (**Fig. 2d**) indicate that COV2-2130 is able to bind RBD in both “up” and “down” conformations of the S trimer. These structural findings are consistent with our previous lower resolution results for the complex using negative stain electron microscopy^1^.

**Fig. 1.**
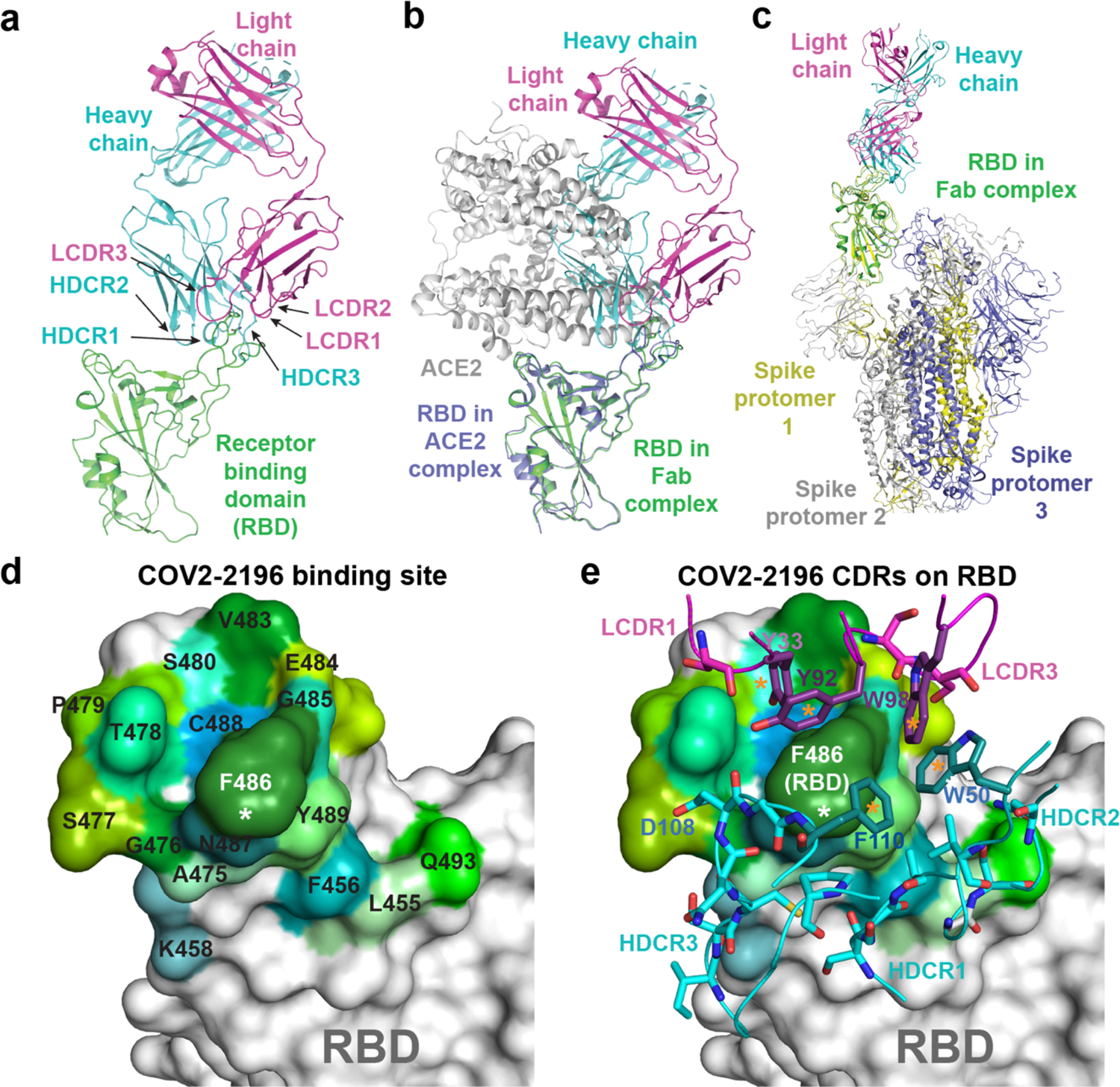
Crystal structure of S protein RBD in complex with Fab COV2-2196. **a.** Cartoon representation of COV2-2196 in complex with RBD. COV2-2196 heavy chain is shown in cyan, light chain in magenta, and RBD in green. **b.** Structure of COV2-2196-RBD complex is superimposed onto the structure of RBD-human ACE2 complex (PDB ID: 6M0J), using the RBD structure as the reference. The color scheme of COV2-2196-RBD complex is the same as that in Fig. 1a. The RBD in the RBD-ACE2 complex is colored in light blue, the human ACE2 peptidase domain in grey. **c.** Structure of COV2-2196-RBD complex is superimposed onto the structure of spike with single RBD in the “up” conformation (PDB ID: 6XM4), using the RBD in “up” conformation as the reference. The color scheme of COV2-2196-RBD complex is the same as that in Fig. 1a. The three subunits of spike are colored in grey, yellow, or light blue respectively (the subunit with its RBD in “up” conformation is yellow). **d.** Surface representation of RBD epitope recognized by COV2-2196. The epitope residues are colored in different shades of green and labeled in black with the critical contact residue F486 labled in white. **e.** Antibody-antigen interactions between COV2-2196 and RBD. RBD is shown in the same surface representation and orientation as that in Fig. 1d. COV2-2196 paratope residues are shown in stick representation. The heavy chain is colored in cyan, and light chain is colored in magenta.

**Fig. 2.**
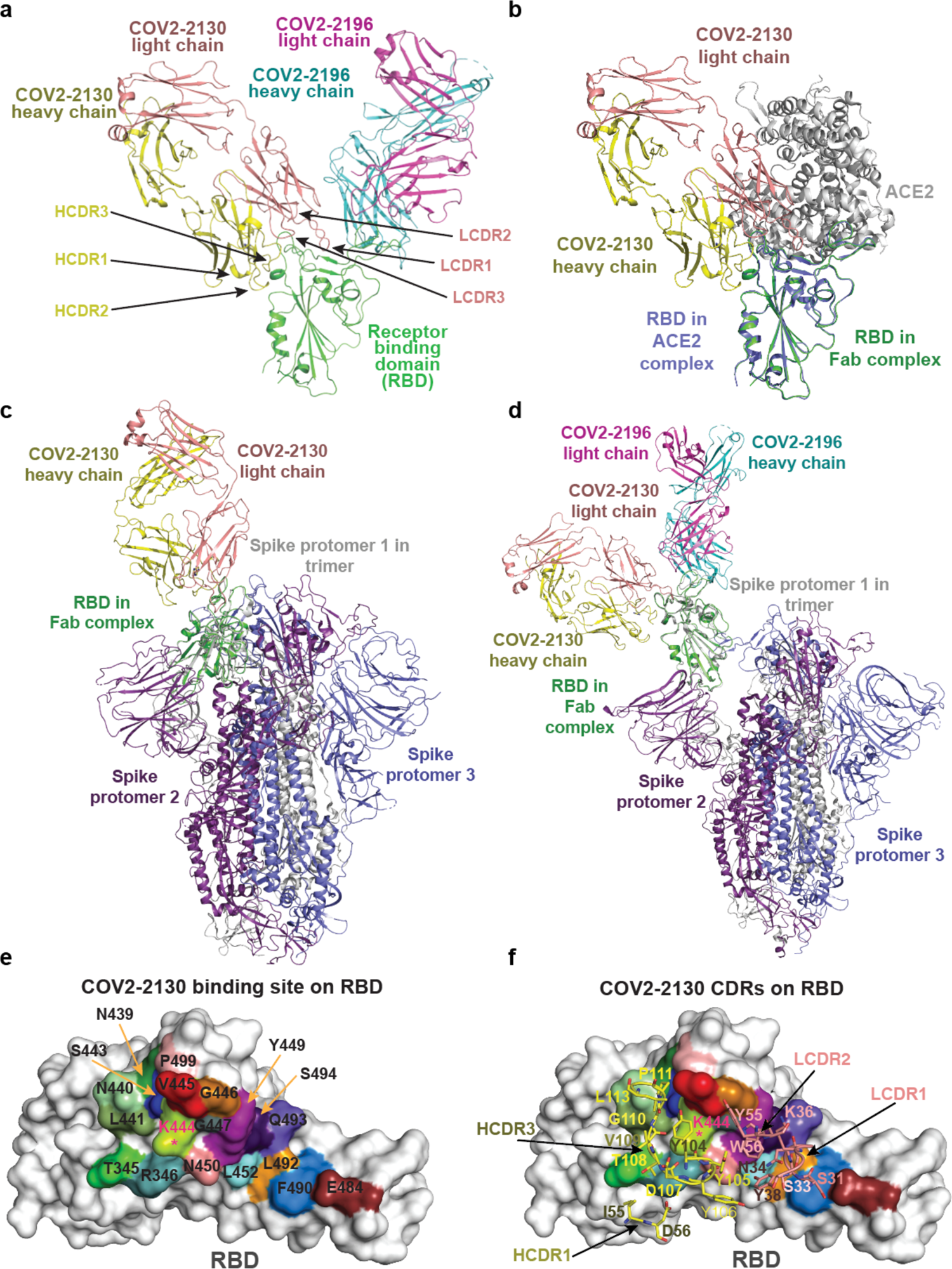
Crystal structure of S protein RBD in complex with both Fabs COV2-2196 and COV2-2130. **a.** Cartoon representation of crystal structure of S protein RBD in complex with COV2-2196 and COV2-2130 Fabs. RBD is shown in green, COV2-2196 heavy chain in cyan, COV2-2196 light chain in magenta, COV2-2130 heavy chain in yellow, and COV2-2130 light chain in orange. CDRs of COV2-2130 are labeled. **b.** Structure of COV2-2130-RBD complex is superimposed onto the structure of the RBD-ACE2 complex (PDB ID: 6M0J), using the RBD structure as the reference. The color scheme of the COV2-2130-RBD complex is the same to that of Fig. 2a. The RBD in the RBD-ACE2 complex is colored in light blue, the human ACE2 peptidase domain in grey. **c.** Structure of COV2-2130-RBD complex is superimposed onto the structure of spike with all RBD in “down” conformation (PDB ID: 6ZOY), using the RBD in one protomer as the reference. The color scheme of COV2-2130-RBD complex is the same as that in Fig. 2a. The three protomers of spike are colored in grey, light blue, or purple respectively. **d.** Structure of COV2-2196 + COV2-2130-RBD complex is superimposed onto the structure of spike with one RBD in “up” conformation (PDB ID: 7CAK), using the RBD in “up” conformation as the reference. The color scheme of COV2-2130-RBD complex is the same as that in Fig. 2a. The three protomers of spike are colored in grey, light blue, or purple respectively. **e.** Surface representation of RBD epitope recognized by COV2-2130. The epitope residues are indicated in different colors and labeled in black. **f.** Interactions of COV2-2130 paratope residues with the epitope. RBD is shown in the same surface representation and orientation as those in Fig. 2e. The paratope residues are shown in stick representation. The heavy chain is colored in yellow, and the light chain in orange.

Structural analysis of COV2-2196 in complex with RBD reveals how COV2-2196 recognizes the receptor-binding ridge on the RBD. One of the major contact residues, F486, situates at the center of the binding site, interacting extensively with the hydrophobic pocket (residue P99 of heavy chain and an “aromatic cage” formed by 5 aromatic side chains) between COV2-2196 heavy/light chains via a hydrophobic effect and van der Waals interactions (**Fig. 1d-e, Extended Data Fig. 2a-b**). A hydrogen bond (H-bond) network, constructed with 4 direct antibody-RBD H-bonds and 16 water-mediated H-bonds, surround residue F486 and strengthen the antibody-RBD interaction (**Extended Data Fig. 2c**). Importantly, for all residues except one (residue P99 of the heavy chain) that interact extensively with the epitope, they are encoded by germline sequences (*IGHV1-58*01* and *IGHJ3*02* for the heavy chain, *IGKV3-20*01* and *IGKJ1*01* for the light chain) (**Fig. 3a**) or only their backbone atoms are involved in the antibody-RBD interactions, such as heavy chain N107 and G99 and light chain S94. We noted another antibody in the literature, S2E12, that is encoded by the same *IGHV/IGHJ* and *IGKV/IGKJ* recombinations, with similar but most likely different *IGHD* genes to those of COV2-2196 (*IGHD2-15* vs *IGHD2-2*)^4^. A comparison of the cryo-EM structure of S2E12 in complex with S protein (PDB 7K4N) suggests that the mAb S2E12 likely uses nearly identical antibody-RBD interactions as those of COV2-2196, although variations in conformations of interface residue side-chains can be seen (**Extended Data Fig. 2d**). For example, the phenyl rings of light chain residue Y92 are perpendicular to each other in the two structures. These analyses suggest that COV2-2196 and S2E12 have similar modes of recognition of RBD.

**Fig. 3.**
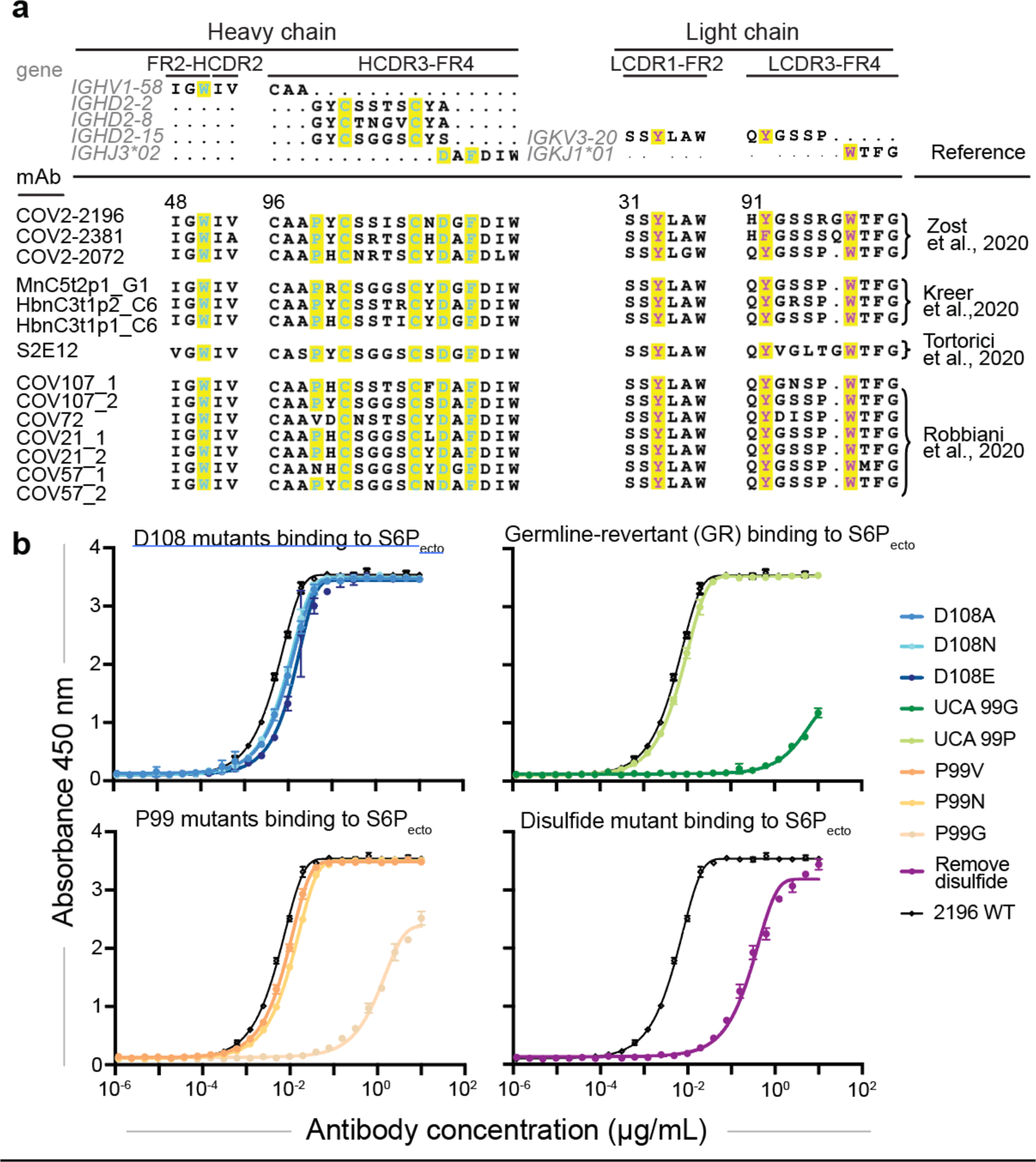
**a.** IMGT/DomainGapAlign results of COV2-2196 heavy and light chains. Key interacting residues and their corresponding residues in germline genes are colored in red. **b.** Binding curves of point mutants of COV2-2196. cDNAs encoding point mutants for the heavy chain, colored in red above, were designed, synthesized as DNA to make recombinant IgG proteins, and tested for binding activity to spike protein. Mutants of D108 residue are in blue, revertant mutation of inferred somatic mutations to germline sequence are in green, P99 mutants are in orange, and a mutant removing the disulfide bond in HCDR3 is in purple.

**Fig. 4.**
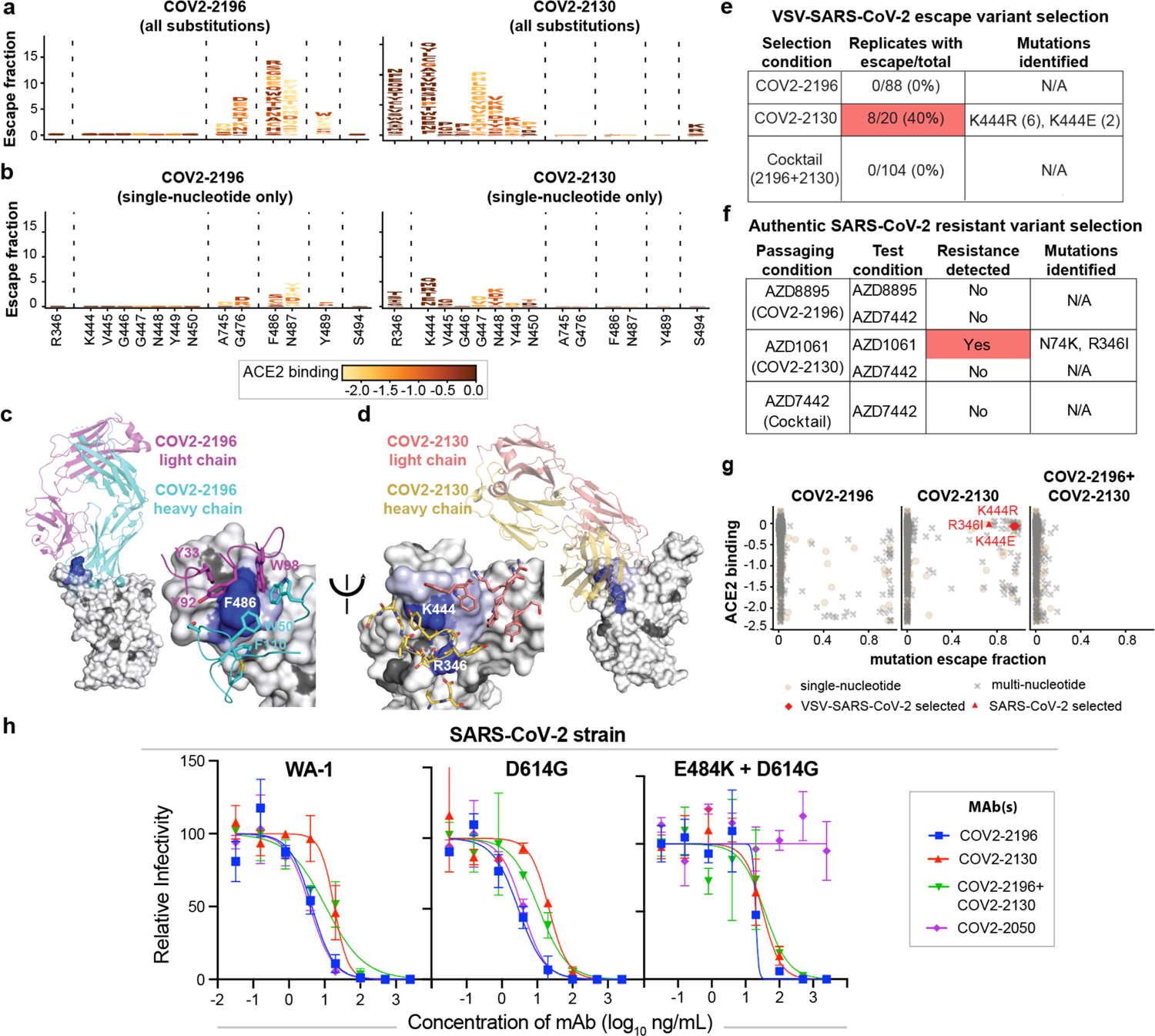
Identification of critical residues for COV2-2196 and COV2-2130 through deep mutational scanning coupled with resistant variant selection. **a.** Logo plots of mutation escape fractions of all at RBD sites with strong escape for COV2-2196 (left) or COV2-2130 (right). Taller letters indicate greater antibody binding escape. Mutations are colored based on the degree to which they reduce RBD binding to human ACE2. Data shown are the average of two independent escape selection experiments using two independent yeast libraries; correlations are shown in **Extended Data** Figure 7b**,c**. Interactive, zoomable versions of these logo plots are at https://jbloomlab.github.io/SARS-CoV-2-RBD_MAP_AZ_Abs/. We determined escape fractions, as described in methods, which represent the estimated fraction of cells expressing that specific variant that fall in the antibody escape bin, such that a value of 0 means the variant is always bound by antibody and a value of 1 means that it always escapes antibody binding. **b.** Logo plots of mutation escape fractions for COV2-2196 and COV2-2130 that are accessible by single nucleotide substitutions from the Wuhan-Hu-1 reference strain used in escape selections (**e,f**). The effect of each substitution on ACE2 binding is represented as in Fig. 4a. **c.** Left panel: mapping deep mutational scanning escape mutations for COV2-2196 onto the RBD surface in the RBD-COV2-2196 structure. Mutations that abrogate COV2-2196 binding are displayed on the RBD structure using a heatmap, where blue represents the RBD site with the greatest cumulative antibody escape and white represents no detected escape. Grey denotes residues where deleterious effects on RBD expression prevented assessment of the effect of the mutation on antibody binding. Right panel: the blow-up of the left panel showing interacting residues around the strongest escape sites of RBD. COV2-2196 heavy chain is colored cyan and the light chain magenta. Two replicates were performed with independent libraries, as described in (a). **d.** Right panel: mapping deep mutational scanning escape mutations for COV2-2130 onto the RBD surface in the RBD-COV2-2130 structure. Mutations that abrogate COV2-2130 binding are displayed on the RBD structure using a heatmap as in Fig. 4c. Left panel: the blow-up of the left panel showing interacting residues around the strongest escape sites of RBD. COV2-2130 heavy chain is colored yellow and the light chain salmon. **e.** Table showing the results of VSV-SARS-CoV-2 escape selection experiments with COV2-2196, COV2-2130, and their combination. The number of escape mutants selected and the total number of escape selection replicates performed is noted, as well as the residues identified by sequencing escape mutant viruses. **f.** Table showing the results of passage of SARS-CoV-2 in the presence of sub-neutralizing concentrations of AZD8895 (based on COV2-2196), AZD1061 (based on COV2-2130), and AZD7442 (AZD8895 + AZD1061). Resistance-associated viral mutations identified by sequencing neutralization-resistant plaques are denoted. **g.** Scatter plot showing DMS data from (**a**), with mutation escape fraction on the x-axis and effect on ACE2 binding on the y-axis. Crosses denote mutations accessible only by multi-nucleotide substitutions, while circles indicate mutations accessible by single-nucleotide substitution. Amino acid substitutions selected by COV2-2130 in VSV-SARS-CoV-2 (K444R, K444E) or authentic SARS-CoV-2 (R346I) are denoted. **h.** Antibody neutralization as measured by FRNT against reference strains and point mutants observed in SARS-CoV-2 variants of concern. Neutralization assays were performed in duplicate and repeated twice, with results shown from one experimental replicate. Error bars denote the range for each point. Mutations compared to the WA-1 reference strain are denoted.

We searched genetic databases to determine if these structural features are present in additional SARS-CoV-2 mAbs isolated by others and found additional members of the clonotype (**Fig 3a**). Two other studies reported the same or a similar clonotype of antibodies isolated from multiple COVID-19 convalescent patients^2, 4^, and one study found three antibodies with the same *IGHV1-58* and *IGKV3-20* pairing, without providing information on D or J gene usage^3^. All of these antibodies are reported to bind SARS-CoV-2 RBD avidly and to neutralize virus with high potency^1–4^. So far, there are only two atomic resolution structures of antibodies encoded by these V_H_(D_H_)J_H_ and V_K_-J_K_ recombinations available, the structure for COV2-2196 presented here and that for S2E12^4^. We performed homology modeling for two additional antibodies of this clonotype from our own panel of anti-SARS-CoV-2 antibodies, designated COV2-2072 and COV2-2381. As expected, given that these antibodies are members of a shared genetic clonotype, the modeled structures of COV2-2072/RBD and COV2-2381/RBD complexes are virtually superimposable with those of COV2-2196/RBD and S2E12/RBD at the antibody-RBD interfaces (**Extended Data Fig. 3a-e**). Additionally, COV2-2072 encodes an N-linked glycosylation sequon in the HCDR3 (**Extended Data Fig. 3d**), an unusual feature for antibodies, given that glycosylation of CDRs might adversely affect antigen recognition. However, the COV2-2196 structure shows that the disulfide-stapled HCDR3 in this clonotype is angled away from the binding site, explaining how this unusual HCDR3 glycosylation in COV2-2072 can be tolerated without compromising binding (**Extended Data Fig. 3e**).

We next determined whether we could identify potential precursors of this public clonotype in the antibody variable gene repertoires of circulating B cells from SARS-CoV-2-naïve individuals. We searched for the V(D)J and VJ genes in previously described comprehensive repertoire datasets originating from 3 healthy human donors, without a history of SARS-CoV-2 infection, and in datasets from cord blood collected prior to the COVID-19 pandemic^26^. A total of 386, 193, 47, or 7 heavy chain sequences for this SARS-CoV-2 reactive public clonotype was found in each donor or cord blood repertoire, respectively (**Extended Data Fig. 4a**). Additionally, we found 516,738 human antibody sequences with the same light chain V-J recombination (*IGKV3-20-IGKJ1*01*). A total of 103,534, 191,039, or 222,165 light chain sequences was found for this public clonotype in each donor respectively. Due to the large number of sequences, the top five abundant sequences were aligned from each donor. Multiple sequence alignments were generated for each donor’s sequences and logo plots were generated. The top 5 sequences with the same recombination event in each donor were identical, resulting in the same logo plots (**Extended Data Fig. 4a-b**).

We noted that 8 of the 9 common residues important for RBD binding in the antibody were encoded by germline gene sequences. Interestingly, these residues were present in all 14 members of the public clonotype that we or others have described (**Fig. 3a**)^1–4^. To validate the importance of these features, we expressed variant antibodies with point mutations in the paratope to determine the effect of variation at conserved residues (**Fig. 3b**).

Altering the D108 residue to A, N, or E had little effect, but removing the disulfide bond in the HCDR3 through cysteine to alanine substitutions greatly reduced binding. While altering the P99 residue to V or N had little effect, a P99G substitution had a dramatic effect on binding. Additionally, we made two germline revertants of the COV2-2196 antibody. The P99 residue is not templated by either the V-gene *IGHV 1-58* nor the D gene *IGHD 2-2*. However, *IGHD 2-2* has a likely templated G at position 99. Therefore, two germline revertants were tested - one with P99 and the other with G99. As the P99 residue orients the HCDR3 loop away from the interaction site with antigen, the G99 germline revertant exhibited reduced binding, whereas the P99 germline revertant bound antigen equivalently to *wt* COV2-2196 (**Fig 3b**).

An antibody based on the COV2-2196 variable region is being tested in combination with an antibody based on the COV2-2130 variable region in clinical trials. Unlike, COV2-2196, COV2-2130 uses the HCDR3 for critical contacts. The HCDR3 comprises 22 amino acid residues, which is relatively long for human antibodies. The HCDR3 forms a long, structured loop that is stabilized by short-ranged hydrogen bonds and hydrophobic interactions/aromatic stackings within the HCDR3, and is further strengthened by its interactions (hydrogen bonds and aromatic stackings) with residues of the light chain (**Extended Data Fig. 5a-b**). The COV2-2130 heavy and light chains are encoded by the germline genes *IGHV3-15* and *IGKV4-1*, respectively, and the two genes encode the longest germline-encoded HCDR2 (10 aa) and LCDR1 (12 aa) loops, which are used in COV2-2130. The heavy chain V(D)J recombination, HCDR3 mutations, and the pairing of heavy and light chains result in a binding cleft between the heavy and light chains, matching the shape of the RBD region centered at S443 – Y449 loop (**Fig. 2a, Extended Data Fig. 5c**). Closely related to these structural features, only HCDR3, LCDR1, HCDR2, and LCDR2 are involved in the formation of the paratope (**Fig. 2e-f, Extended Data Fig. 2e-f**). Inspection of the antibody-RBD interface reveals a region that likely drives much of the energy of interaction. The RBD residue K444 sidechain is surrounded by subloop Y104 – V109 of the HCDR3 loop, and the positive charge on the side chain nitrogen atom is neutralized by the HCDR3 residue D107 side chain, three mainchain carbonyl oxygen atoms from Y105, D107, and V109, and the electron-rich face of the Y104 phenyl ring (cation-π interaction) (**Extended Data Fig. 2e**). Since the interacting atoms are completely protected from solvent, the highly concentrated interactions within such a restricted space are energetically favorable. Furthermore, this “hotspot” of the antibody-RBD interface is surrounded by or protected from the solvent by antibody-RBD interactions with lesser free energy gains, including salt bridge between the RBD residue R346 and HCDR2 D56, electrostatic interaction between RBD R346 and the mainchain oxygen of HCDR3 Y106, a hydrogen bond between RBD N450 and HCDR3 Y105 mainchain oxygen, a hydrogen bond between RBD V445 mainchain oxygen and HCDR3 Y104 sidechain, a hydrophobic interaction between V445 sidechain and sidechains of HCDR3 L113 and F118 (**Extended Data Fig. 2e**). Also, aromatic stacking between the HCDR3 residue Y105 and LCDR2 residue W56 participates in the shielding of the “hotspot” from solvent (**Extended Data Fig. 2e**). In addition, COV2-2130 light chain LCDR1 and LCDR2 make extensive contacts with the RBD. Among them, the LCDR1 S32 sidechain, S33 mainchain oxygen, N34 sidechain, and LCDR2 Y55 sidechain form hydrogen bonds with RBD E484 sidechain, S494 mainchain nitrogen, Y449 mainchain oxygen, and G446 mainchain nitrogen (**Extended Data Fig. 2f**). Residues LCDR1 K36, Y38, and LCDR2 W56 interact with the RBD Y449 via aromatic stackings and cation-π interactions, forming an “interaction cluster” (**Extended Data Fig. 2f**), although these interactions are likely not energetically as strong as in the case of RBD K444. In the crystal structure of the RBD in complex with both COV2-2196 and COV2-2130, we noted a possible interaction between the closely spaced COV2-2196 and COV2-2130 Fabs (**Extended Data Fig. 6**).

To better understand the RBD sites critical for binding of COV2-2196 and COV2-2130, we used a deep mutational scanning (DMS) approach to map all mutations to the RBD that escape antibody binding^27^; (**Extended Data Fig. 7**). For both antibodies, we identified several key positions, all in the antibody binding site, where RBD mutations strongly disrupted binding (**Fig. 4a-d**). We leveraged our previous work quantifying the effects of RBD mutations on ACE2 binding^28^ to overlay the effect on ACE2 binding for mutations that abrogated antibody binding to RBD (**Fig. 4a,b)**. For COV2-2196, many mutations to F486 and N487 had escape fractions approaching 1 (*i.e*., those RBD variants to which the antibody does not bind), reinforcing the importance of the contributions of these two residues to antibody binding. Similarly, for COV2-2130, mutation of residue K444 to any of the other 19 amino acids abrogated antibody binding, indicating that the lysine at this position is critical for the antibody-RBD interaction.

Nevertheless, not all antibody binding site residues were identified as sites where mutations greatly reduced binding. Several explanations are possible: 1) some binding site residues may not be critical for binding, 2) some RBD residues do not use their side chains to form interactions with the mAbs or 3) mutations at some sites may not be tolerated for RBD expression^28^. For instance, residues L455, F456, and Q493 are part of the structurally-defined binding site for COV2-2196 (**Fig. 1d)**, but mutations to these sites did not impact antibody binding detectably (**Fig. 4a and c**), suggesting that these residues do not make critical binding contributions. Superimposition of the COV2-2196/RBD structure onto the S2E12/RBD structure clearly demonstrates a flexible hinge region between the RBD ridge and the rest of the RBD that is maintained when antibody is bound (**Extended Data Fig. 2d**). This finding indicates that mutations at these three positions could be well-tolerated for antibody-RBD binding and supports the non-essential nature of these particular residues for COV2-2196 or S2E12 binding.

Importantly, COV2-2196 and COV2-2130 do not compete with one another for binding to the RBD^1^, suggesting they could comprise an escape-resistant cocktail for prophylactic or therapeutic use. Indeed, the binding sites and escape variant maps for these two antibodies are non-overlapping. To test whether there were single mutations that could escape binding of both antibodies, we performed escape variant mapping experiments with a 1:1 mixture of the COV2-2196 and COV2-2130 antibodies, but we did not detect any mutation that had an escape fraction of greater than 0.2, whereas the mutations with the largest effects for each of the single antibodies was approximately 1 (**Extended Data Fig. 7d**).

Although these experiments map all mutations that escape antibody binding to the RBD, we also sought to determine which mutations have the potential to arise during viral growth. To address this question, we first attempted to select escape mutations using a recombinant VSV expressing the SARS-CoV-2 S glycoprotein (VSV-SARS-CoV-2)^29^; (**Fig 4e**). We expected that the only amino acid mutations that would be selected during viral growth were those 1) arising by single-nucleotide RNA changes, 2) causing minimal deleterious effect on ACE2 binding and expression, and 3) substantially impacting antibody binding^27, 28^. Indeed, we did not detect any COV2-2196-induced mutations that were both single-nucleotide accessible and relatively well-tolerated with respect to effects on ACE2 binding (**Fig. 4b**), which may explain why escape mutants were not selected in any of the 88 independent replicates of recombinant VSV growth in the presence of antibody (**Fig. 4e Extended Data Fig. 7g**). For COV2-2130, mutations to site K444, a site that is relatively tolerant to mutation^28^, demonstrated the most frequent escape from antibody binding in neutralization assays with the the VSV chimeric virus. K444R (selected in 6 out of 20 replicates) or K444E (selected in 2 out of 20 replicates) were identified in 40% of the replicates of recombinant VSV growth in the presence of COV2-2130 (**Fig. 4e, Extended Data Fig. 7g**).

To explore resistance with authentic infectious virus, SARS-CoV-2 strain USA-WA1/2020 was passaged serially in Vero cell monolayer cultures with the clinical antibodies based on COV2-2196 (AZD8895), COV2-2130 (AZD1061) or their 1:1 combination (AZD7442), at concentrations beginning at their respective IC_50_ values and increased step-wise to their IC_90_ value with each passage (**Extended Data Fig. 8)**. As a control, virus was passaged in the absence of antibody. Following the final passage, viruses were evaluated for susceptibility against the partner antibody at a final concentration of 10 times the IC_90_ concentration by plaque assay. We did not detect any plaques resistant to neutralization by AZD8895 (based on COV2-2196) or the AZD7442 cocktail. Virus that was passaged serially in AZD1061 formed plaques to a titer of 1.2 × 10^7^ PFU/mL after selection in 10 × the IC_90_ value concentration of AZD1061, but plaques were not formed with AZD7442. Plaques (n=6) were selected randomly, and the S gene was amplified and sequenced, revealing the same 3 amino acid changes in all 6 of the independently selected and sequenced plaques: N74K, R346I and S686G (**Fig. 4f**). The S686G change has been reported previously to be associated with serial passaging of SARS-CoV-2 in Vero cells^30^, isolated from challenge studies in ferrets^31^ or NHPs^32^, and is predicted to decrease furin activity^30^. The N74K residue is located in the N-terminal domain outside of the AZD1061 binding site and results in the loss of a glycan^33^. The R346I residue is located in the binding site of AZD1061 and may be associated with AZD1061-resistance. The impact of the R346I changes on AZD1061 (COV2-2130) binding to S protein is shown in **Fig. 4g**. The K444R and K444E substitutions selected in the VSV-SARS-CoV-2 system and the R346I substitution selected by passage with authentic SARS-CoV-2 are accessible by single nucleotide substitution and preserve ACE2 binding activity (**Fig. 4g**), indicating that our DMS analysis predicted the mutations selected in the presence of COV2-2130 antibody. Taken together, these results comprehensively map the effects of all amino acid substitutions on the binding of COV2-2196 and COV2-2130 and identify sites of possible concern for viral evolution. That said, variants containing mutations at residues K444 and R346 are rare among all sequenced viruses present in the GISAID databases (all ≤ 0.01% when accessed on 12/23/20).

Recently, viral variants with increased transmissibility and potential antigenic mutations have been reported in clinical isolates^34–37^. We tested whether some of the variant residues in these rapidly emerging strains would abrogate the activity of these potently neutralizing antibodies. We tested isogenic D614G and E484K variants in the WA-1 strain background (2019n-CoV/USA_WA1/2020, [WA-1]), all prepared as authentic SARS-CoV-2 viruses and used in focus reduction neutralization tests^29^. The E484K mutation was of special interest, since this residue is located within 5Å of each of the mAbs in the complex of Fabs and RBD, albeit at the very binding site. E484K also is present in emerging lineages B.1.351 (501Y.V2)^36^ and P.1 (501Y.V3)^37^, and has been demonstrated to alter the binding of some monoclonal antibodies^38, 39^ as well as human polyclonal serum antibodies^40^. Variants containing E484K also have been shown to be neutralized less efficiently by convalescent serum and plasma from SARS-CoV-2 survivors^41–43^. For COV2-2196, COV2-2130, and COV2-2050 (a third neutralizing antibody we incuded for comparison as it interacts with the residue E484), we found virtually no impact of the D614G mutation (**Fig. 4h**). However, we did observe effects on neutralization with the D614G/E484K virus. COV2-2050 completely lost neutralization activity, consistent with our previous study defining E484K as a mutation abrogating COV2-2050 binding^27^. In contrast, COV2-2196, COV2-2130, and COV2-2196 + COV2-2130 showed only minor reduction in inhibitory capacity (2- to 5-fold increases in IC_50_ values). Recent reports from others with neutralization data for recombinant forms of COV2-2196 and COV2-2130 against a viral variant containing all the RBD substitutions in the Sotuh African lineage, rather than just the E484K substitution, also show that this antibody cocktail is effective against emerging variants of concern^44–47^.

## Discussion

The process of B cell development, in which diverse variable gene segments are recombined, results in human naïve B cell repertoires containing an enormous amount of structural diversity in the complementarity determining regions (CDRs) of the antibodies (Abs) that they encode. Despite this extensive and diverse pool of naïve B cells, infection or vaccination with viral pathogens sometimes elicit antibodies in diverse individuals that share common structural features encoded by the same antibody variable genes. Examples of recurring variable gene usage have been described for antibody responses to human rotavirus^21, 48^, human immunodeficiency virus^49–52^ influenza A virus^53–56^, and hepatitis C virus^57, 58^, among others. The recognition of the use of common variable genes in antiviral responses has led to the general concept of B cell public clonotypes, or B cells with similar genetic features in their variable regions that encode for antibodies with similar patterns of specificity and function in different individuals. A number of recent reports have described the identification of public clonotypes in the Ab responses to SARS-CoV-2^2, 14, 59, 60^. Identifying and understanding the genetic and structural basis for selection of public clonotypes is valuable, as this information forms the central conceptual underpinning for many current rational structure-based vaccine design efforts^61^. Our structural analyses define the molecular basis for the frequent selection of a public clonotype of human antibodies sharing heavy chain V-D-J and light chain V-J recombinations that target the same region of the SARS-CoV-2 S RBD. Germline antibody gene-encoded residues in heavy and light chains play a vital role in antigen recognition, suggesting that few somatic mutations are required for antibody maturation of this clonotype. The existence of potenty neutralizing public clonotypes across multiple individuals may in part account for the remarkable efficacy of S protein-based vaccines that is being observed in the clinic. One might envision an opportunity to elicit serum neutralizing antibody titers with even higher neutralization potency using domain- or motif-based vaccine designs for this antigenic site to prime human immune responses to elicit this clonotype.

The recent emergence of variant virus lineages with increased transmissibility and altered sequences in known sites of neutralization is concerning for the capacity of SARS-CoV-2 to evade current antibody countermeasures in development and testing. Our comprehensive mapping of the effect of RBD mutations on the binding of COV2-2196 and COV2130 underscores their use as a rationally designed cocktail, given that they have orthogonal escape mutations. Our DMS experiments are also consistent with the binding site determined by our antibody-RBD crystal structures and the DMS results predict the mutations present in resistant variants selected by *in vitro* passaging experiments. We tested the activity of the individual antibodies or the cocktail against recombinant authentic viruses containing mutations from several important variants of concern, and demonstrate that the individual antibodies or their combination are capable of potently neutralizing these emerging variants. Recent work from others also has demonstrated that some circulating variants of concern exhibit substantial escape from neutralization of many human monoclonal antibodies in clinical development, but recombinant forms of COV2-2196 and COV2-2130 still potently neutralized pseudoviruses that included the emerging B.1.1.7 and B.1.351 lineages^44^. Taken together, this work defines the molecular basis for potent neutralization of SARS-CoV-2 by COV2-2196 and COV2-2130 and demonstrates that these antibodies efficiently neutralize emerging antigenic variants either separately or in combination, underscoring the promise of the AZD7442 investigational cocktail for use in the prevention and treatment of COVID-19.

### Data and materials availability

The crystal structures reported in this paper have been deposited to the Protein Data Bank (https://www.rcsb.org) under the accession numbers 7L7D (COV2-2196 + RBD) and 7L7E COV2-2196 and COV2-2130 + RBD). The following were obtained from the PDB and used for visualization or molecular replacement: PDB IDs: 7K4N, 6M0J, 6XM4, 7CAK, 6ZOY, 6XC2, 5JRP. Sequence Read Archive deposition for the aligned human antibody gene repertoire data set is deposited at the NCBI: PRJNA511481. All other data are available in the main text or the supplementary materials. Requests for reagents may be directed to and be fulfilled by the Lead Contact: Dr. James E. Crowe, Jr. Materials reported in this study will be made available but may require execution of a Materials Transfer Agreement.

### Software availability

The computational pipeline for the deep mutational scanning analysis of antibody escape mutations is available on GitHub: https://github.com/jbloomlab/SARS-CoV-2-RBD_MAP_AZ_Abs. The FASTQ files are available on the NCBI Sequence Read Archive under BioSample SAMN17532001 as part of BioProject PRJNA639956.. Per-mutation escape fractions are available on GitHub (https://github.com/jbloomlab/SARS-CoV-2-RBD_MAP_AZ_Abs/blob/main/results/supp_data/AZ_cocktail_raw_data.csv) and in **Supplementary Data Table 1**.

## Acknowledgments

At Fred Hutchinson Cancer Research Center, we thank Amin Addetia for experimental assistance, the Flow Cytometry and Genomics core facilities, and Scientific Computing, supported by ORIP grant S10OD028685. We thank Adrian Creanga and Barney Graham of the U.S. National Institutes of Health (N.I.H.) for the Vero-hACE2-TMPRSS2 cells. At AstraZeneca, we thank Paul Warrener, Christopher Morehouse and Dave Tabor for virus genome sequencing and spike variant analysis, and Kuishu Ren for generation of protein reagents and related binding data.

## Author contributions

Conceptualization, J.D., S.J.Z., J.D.B. and J.E.C.; Investigation, J.D., S.J.Z., A.J.G., T.N.S, A.S.D., E.C.C., R.E.C., J.B.C., R.E.S., P.G., J.R., E.A., C.G., R.S.N.; E.B., X.X., X.Z., J.L., S.W., M.E.M., M.B.F., T.B., K.T., H.B., Y.M-L., P.M.; Writing – Original Draft, J.D., S.J.Z. and J.E.C; All authors edited the manuscript and approved the final submission); Supervision, P.-Y.S, M.E., M.S.D., J.D.B., J.E.C.; Funding acquisition, P.-Y.S., M.E., R.H.C., M.SD., J.D.B., J.E.C.

## Additional information

**Supplementary information** is available for this paper.

Correspondence and requests for materials should be addressed to J.E.C.

---

**Extended Data Fig. 1. Overlay of substructure of RBD-COV2-2196 in RBD-COV2-2196-2130 complex and RBD-COV2-2196 crystal structure.**

**Extended Data Fig. 2. Similar aromatic stacking and hydrophobic interaction patterns at the RBD site F486 shared between RBD-COV2-2196 and spike-S2E12 complexes.**

**a.** Same hydrogen bonding pattern surrounding residue F486 in the structures of the two complexes.

**b.** Detailed interactions between COV2-2196 and RBD. COV2-2196 heavy chain is colored in cyan, the light chain is colored in magenta, and RBD is colored in green. Important interacting residues are shown in stick representation. Water molecules involved in antibody-RBD interaction are represented as pink spheres. Direct hydrogen bonds are shown as orange dashed lines, and water-mediated hydrogen bonds as yellow dashed lines.

**c.** Superimposition of S2E12/RBD cryo-EM structure onto the COV2-2196/RBD crystal structure, with the variable domains of antibodies as references. COV2-2196 heavy chain is in cyan, and its light chain in magenta; S2E12 heavy chain is in pale cyan, and its light chain in light pink. The two corresponding RBD structures are colored in green or yellow, respectively.

**d.** Detailed interactions between COV2-2130 heavy chain and RBD. Paratope residues are shown in stick representation and colored in yellow, epitope residues in green sticks. Hydrogen-bonds or strong polar interactions are represented as dashed magenta lines.

**e.** Detailed interactions between COV2-2130 light chain and RBD. Paratope residues are shown in stick representation and colored in orange, epitope residues in green sticks. Hydrogen-bonds are represented as dashed magenta lines.

**Extended Data Fig. 3. A common clonotype of anti-RBD antibodies with the same binding mechanism.**

**a.** COV2-2196/RBD crystal structure.

**b.** S2E12/RBD cryo-EM structure.

**c.** COV2-2381/RBD homology model. COV2-2072 encodes an N-linked glycosylation sequon in the HCDR3, indicated by the gray spheres.

**d.** COV2-2072/RBD homology model.

**e.** Overlay of the COV2-2196/RBD crystal structure (**a**) and S2E12/RBD cryo-EM structure (**b**).

**Extended Data Fig. 4. Identification of putative public clonotype members genetically similar to COV2-2196 in the antibody variable gene repertoires of virus-naïve individuals.**

Antibody variable gene sequences collected from healthy individuals prior to the pandemic with the same sequence features as COV2-2196 heavy chain (a) and light chain (b) are aligned. Sequences from three different donors as well as cord blood included sequences with the features of the public clonotype. The sequence features and contact residues used in COV2-2196 are highlighted in red boxes below each multiple sequence alignment.

**Extended Data Fig. 5.**

**a.** Detailed COV2-2130 HCDR3 loop structure. Short-range hydrogen bonds, stabilizing the loop conformation, are shown as dashed magenta lines.

**b.** Residues of COV2-2130 light chain form aromatic stacking interactions and hydrogen bonds with HCDR3 to further stabilize the HCDR3 loop.

**c.** Long LCDR1, HCDR2, and HCDR3 form complementary binding surface to the RBD epitope. RBD is shown as surface representation in grey. COV2-2130 heavy chain is colored in yellow with HCDR3 in orange, and the light chain in salmon with LCDR1 in magenta.

**d.** 180° rotation view of panel **c**.

**Extended Data Fig. 6. Interface between COV2-2196 and COV2-2130 in the crystal structure of RBD in complex with COV2-2196 and COV2-2130.**

COV2-2196 heavy or light chain are shown as cartoon representation in cyan or magenta, respectively, and COV2-2130 heavy or light chain in yellow or salmon, respectively. The RBD is colored in green. Interface residues are shown in stick representation.

**Extended Data Fig. 7. Identification by deep mutational scanning of mutations affecting antibody binding and method of selection of antibody resistant mutants with VSV-SARS-CoV-2 virus.**

**a.** Top: Flow cytometry plots showing representative gating strategy for selection of single yeast cells using forward- and side-scatter (first three panels) and selection of yeast cells expressing RBD (right panel). Each plot is derived from the preceding gate. Bottom: Flow cytometry plots showing gating for RBD^+^, antibody^-^ yeast cells (*i.e.*, cells that express RBD but where a mutation prevents antibody binding). Selection experiments are shown for COV2-2196 or COV2-2130, with two independent libraries shown for each.

**b.** Correlation of observed sites of escape from antibody binding between yeast library selection experiments using COV2-2196, COV2-2130, or a 1:1 mixture of COV2-2196 and COV2-2130. The x-axes show cumulative escape fraction for each site for library 1, and the y-axes show cumulative escape fraction for each site for library 2. Correlation coefficient and *n* are denoted for each graph.

**c.** Correlation of observed mutations that escape antibody binding between yeast library selection experiments using COV2-2196, COV2-2130, or a 1:1 mixture of COV2-2196 and COV2-2130. The x-axes show each amino acid mutation’s escape fraction for library 1, and the y-axes show each amino acid mutation’s escape fraction for library 2. Correlation coefficient and *n* are denoted for each graph.

**d-f.** DMS results for COV2-2196 (**d**), COV2-2130 (**e**), or a 1:1 mixture of COV2-2196 and COV2 2130 (**f**). Left panels: sites of escape across the entire RBD are indicated by peaks that correspond to the logo plots in the middle and right panel. Middle panel: as in Fig. 4a, logo plot of cumulative escape mutation fractions of all RBD sites with strong escape mutations for COV2-2196, or COV2-2130, or COV2-2196+COV2-2130. Mutations are colored based on the degree to which they abrogate RBD binding to human ACE2. Right panel: again, logo plots show cumulative escape fractions, but colored based on the degree to which mutations effect RBD expression in the yeast display system. Interactive, zoomable versions of these logo plots are at https://jbloomlab.github.io/SARS-CoV-2-RBD_MAP_AZ_Abs/.

**g.** Representative RTCA sensograms showing virus that escaped antibody neutralization. Cytopathic effect (CPE) was monitored kinetically in Vero E6 cells inoculated with virus in the presence of a saturating concentration (5 μg/mL) of antibody COV2-2130. Representative instances of escape (magenta) or lack of detectable escape (blue) are shown. Uninfected cells (green) or cells inoculated with virus without antibody (red) serve as controls. Magenta and blue curves represent a single representative well; the red and green controls are the mean of technical duplicates.

**h.** Representative RTCA sensograms validating that a variant virus selected by COV2-2130 in (**g**) indeed escaped COV2-2130 (magenta) but was neutralized by COV2-2196 (light blue).

**i.** Example sensograms from individual wells of 96-well E-plate analysis for escape selection experimetnts with COV2-2196, COV2-2130, or a 1:1 mix of COV2-2196 and COV2-2130. Instances of escape from COV2-2130 are noted, while escape was not detected in the presence of COV2-2196 or COV2-2196+COV2-2130. Positive and negative controls are denoted on the first plate.

**Extended Data Fig. 8. Method of selection of antibody resistant mutants with authentic SARS-CoV-2 virus.**

The method for assessing monoclonal antibody resistant spike protein variants is shown. SARS-CoV-2 was passaged serially in the presence of monoclonal antibodies at the increasing concentrations indicated in the figure or without antibody (no monoclonal antibody). Following passage at IC_90_ concentrations, samples were treated with 10 × IC_90_ concentrations of monoclonal antibodies and any resultant resistant virus collected, and the genome was sequenced.

## Materials and Methods

### Expression and purification of recombinant receptor binding domain (RBD) of SARS-CoV-2 spike protein

The DNA segments correspondent to the S protein RBD (residues 319 - 528) was sequence optimized for expression, synthesized, and cloned into the pTwist-CMV expression DNA plasmid downstream of the IL-2 signal peptide (MYRMQLLSCIALSLALVTNS) (Twist Bioscience). A three amino acid linker (GSG) and a His-tag were incorporated at the C-terminus of the expression constructs to facilitate protein purification. Expi293F cells were transfected transiently with the plasmid encoding RBD, and culture supernatants were harvested after 5 days. RBD was purified from the supernatants by nickel affinity chromatography with HisTrap Excel columns (GE Healthcare Life Sciences). For protein production used in crystallization trials, 5 μM kifunensine was included in the culture medium to produce RBD with high mannose glycans. The high mannose glycoproteins subsequently were treated with endoglycosidase F1 (Millipore) to obtain homogeneously deglycosylated RBD.

### Expression and purification of recombinant COV2-2196 and COV2-2130 Fabs

The DNA fragments corresponding to the COV2-2196 and COV2-2130 heavy chain variable domains with human IgG1 CH1 domain and light chain variable domains with human kappa chain constant domain were synthesized and cloned into the pTwist vector (Twist Bioscience). This vector includes the heavy chain of each Fab, followed by a GGGGS linker, a furin cleavage site, a T2A ribosomal cleavage site, and the light chain of each Fab. Expression of the heavy and light chain are driven by the same CMV promoter. COV2-2196 and COV2-2130 Fabs were expressed in ExpiCHO cells by transient transfection with the expression plasmid. The recombinant Fab was purified from culture supernatant using an anti-CH1 CaptureSelect column (Thermo Fisher Scientific). For the RBD/COV2-2196 complex, the *wt* sequence of COV2-2196 was used for expression. For the RBD/COV2-2196/COV2-2130 complex, a modified version of COV2-2196 Fab was used in which the first two amino acids of the variable region were mutated from QM to EV.

### Crystallization and structural determination of antibody-antigen complexes

Purified COV2-2196 Fab was mixed with deglycosylated RBD in a molar ratio of 1:1.5, and the mixture was purified further by size-exclusion chromatography with a Superdex-200 Increase column (GE Healthcare Life Sciences) to obtain the antibody-antigen complex. To obtain RBD/COV2-2196/COV2-2130 triple complex, purified and deglycosylated RBD was mixed with both COV2-2196 and COV2-2130 Fabs in a molar ratio of 1:1.5:1.5, and the triple complex was purified with a Superdex-200 Increase column. The complexes were concentrated to about 10 mg/mL and subjected to crystallization trials. The RBD/COV2-2196 complex was crystallized in 16% −18% PEG 3350, 0.2 Tris-HCl pH 8.0 – 8.5, and the RBD/COV2-2196/COV2-2130 complex was crystallized in 5% (w/v) PEG 1000, 100 mM sodium phosphate dibasic/citric acid pH 4.2, 40% (v/v) reagent alcohol. Cryo-protection solution was made by mixing crystallization solution with 100% glycerol in a volume ratio of 20:7 for crystals of both complexes. Protein crystals were flash-frozen in liquid nitrogen after a quick soaking in the cryo-protection solution. Diffraction data were collected at 100 K at the beamline 21-ID-F (wavelength: 0.97872 Å) for RBD/COV2-2196 complex and 21-ID-G (wavelength: 0.97857 Å) for RBD/COV2-2196/COV2-2130 complex at the Advanced Photon Source. The diffraction data were processed with XDS^62^ and CCP4 suite^63^. The crystal structures were solved by molecular replacement using the structure of RBD in complex with Fab CC12.1 (PDB ID 6XC2) and Fab structure of MR78 (PDB ID 5JRP) with the program Phaser^64^. The structures were refined and rebuilt manually with Phenix^65^ or Coot^66^, respectively. The Ramachandran statistics for final structure of RBD-COV2-2196 are: 95.82% favored, 4.18% allowed, and 0.00% disallowed, and the Ramachandran statistics for final structure of RBD-COV2-2196-2130: 95.34% favored, 4.37% allowed, and 0.00% disallowed. The models have been deposited into the Protein Data Bank. PyMOL software^67^ was used to make all of the structural figures.

### COV2-2196 mutant generation

Struturally-important residues in the COV2-2196 heavy chain sequence were identified as D108, P99, and the disulfide bond in HCDR3. The D108 residue was mutated to alanine, asparagine, and glutamic acid. The P99 residue was mutanted to valine, asparagine, and glycine. The disulfide bond was removed by replacing the cystines with alanine. Additionally, the germline revertant forms of COV2-2196 were generated by aligning the sequence to identified germline sequences using IgBlast, and reverting back the residues that were not germline-encoded. DNA fragments corresponding to the COV2-2196 mutant heavy chain variable domains with human IgG1 and light chain variable domain with human kappa chain constant domain were synthesized and cloned into the pTwist_mCis vector (Twist Bioscience) as previously described^25^. Constructs were transformed into *E. coli*, and DNA was purified. Antibodies then were produced by transient tranfection of ExpiCHO cells following the manufacturer’s protocol (Gibco). Supernatants were filter-sterilized using 0.45 µm pore size filters and samples were applied to HiTrap MabSelect Sure columns (Cytiva).

### ELISA binding of COV2-2196 mutants

Wells of 384-well microtiter plates were coated with purified recombinant SARS-CoV-2 S 6P protein at 4°C overnight. Plates were blocked with 2% non-fat dry milk and 2% normal goat serum in DPBS containing 0.05% Tween-20 (DPBS-T) for 1 h. Antibodies were diluted to 10 µg/mL and titrated two-fold 23 times in DPBS-T and added to the wells, followed by an incubation for 1 h at room temperature. The bound antibodies were detected using goat anti-human IgG conjugated with horseradish peroxidase (Southern Biotech) and TMB substrate (Thermo Fischer Scientific). Reactions were quenched with 1 N hydrochloric acid and absorbance was measured at 450 nm using a spectrophotometer (Biotek).

### Mapping of all mutations that escape antibody binding

All mutations that escape antibody binding were mapped via a DMS approach^27^. We used previously described yeast-display RBD mutant libraries^27, 28^. Briefly, duplicate mutant libraries were constructed in the spike receptor binding domain (RBD) from SARS-CoV-2 (isolate Wuhan-Hu-1, Genbank accession number MN908947, residues N331-T531) and contain 3,804 of the 3,819 possible amino-acid mutations, with >95% present as single mutants. Each RBD variant was linked to a unique 16-nucleotide barcode sequence to facilitate downstream sequencing. As previously described, libraries were sorted for RBD expression and ACE2 binding to eliminate RBD variants that are completely misfolded or non-functional (*i.e*., lacking modest ACE2 binding affinity^27^).

Antibody escape mapping experiments were performed in biological duplicate using two independent mutant RBD libraries, as previously described^27^, with minor modifications. Briefly, mutant yeast libraries induced to express RBD were washed and incubated with antibody at 400 ng/mL for 1 h at room temperature with gentle agitation. After the antibody incubations, the libraries were secondarily labeled with 1:100 FITC-conjugated anti-MYC antibody (Immunology Consultants Lab, CYMC-45F) to label for RBD expression and 1:200 PE-conjugated goat anti-human-IgG (Jackson ImmunoResearch 109-115-098) to label for bound antibody. Flow cytometric sorting was used to enrich for cells expressing RBD variants with reduced antibody binding via a selection gate drawn to capture unmutated SARS-CoV-2 cells labeled at 1% the antibody concentration of the library samples. For each sample, approximately 10 million RBD+ cells were processed on the cytometer. Antibody-escaped cells were grown overnight in SD-CAA (6.7 g/L Yeast Nitrogen Base, 5.0 g/L Casamino acids, 1.065 g/L MES acid, and 2% w/v dextrose) to expand cells prior to plasmid extraction.

Plasmid samples were prepared from pre-selection and overnight cultures of antibody-escaped cells (Zymoprep Yeast Plasmid Miniprep II) as previously described^27^. The 16-nucleotide barcode sequences identifying each RBD variant were amplified by PCR and sequenced on an Illumina HiSeq 2500 with 50 bp single-end reads as described^27, 28^.

Escape fractions were computed as described^27^, with minor modifications as noted below. We used the dms_variants package (https://jbloomlab.github.io/dms_variants/, version 0.8.2) to process Illumina sequences into counts of each barcoded RBD variant in each pre-sort and antibody-escape population using the barcode/RBD look-up table previously described^68^.

For each antibody selection, we computed the “escape fraction” for each barcoded variant using the deep sequencing counts for each variant in the original and antibody-escape populations and the total fraction of the library that escaped antibody binding via a previously described formula^27^. These escape fractions represent the estimated fraction of cells expressing that specific variant that fall in the antibody escape bin, such that a value of 0 means the variant is always bound by serum and a value of 1 means that it always escapes antibody binding. We then applied a computational filter to remove variants with low sequencing counts or highly deleterious mutations that might cause antibody escape simply by leading to poor expression of properly folded RBD on the yeast cell surface^27, 28^. Specifically, we removed variants that had (or contained mutations with) ACE2 binding scores < −2.35 or expression scores < −1, using the variant- and mutation-level deep mutational scanning scores as previously described^28^. Note that these filtering criteria are slightly more stringent than those previously used to map a panel of human antibodies^27^ but are identical to those used in recent studies defining RBD residues that impact the binding of mAbs^68^ and polyclonal serum^40^.

We next deconvolved variant-level escape scores into escape fraction estimates for single mutations using global epistasis models^69^ implemented in the dms_variants package, as detailed at (https://jbloomlab.github.io/dms_variants/dms_variants.globalepistasis.html) and described^27^. The reported escape fractions throughout the paper are the average across the libraries (correlations shown in **Extended Data Fig. 7a,b**); these scores are also in **Supplementary Data Table 1.** Sites of strong escape from each antibody for highlighting in logo plots were determined heuristically as sites whose summed mutational escape scores were at least 10 times the median sitewise sum of selection, and within 10-fold of the sitewise sum of the most strongly selected site. Full documentation of the computational analysis is at https://github.com/jbloomlab/SARS-CoV-2-RBD_MAP_AZ_Abs. These results are also available in an interactive form at https://jbloomlab.github.io/SARS-CoV-2-RBD_MAP_AZ_Abs/.

### Antibody escape selection experiments with VSV-SARS-CoV-2

For escape selection experiments with COV2-2196 and COV2-2130, we used a replication competent recombinant VSV virus encoding the spike protein from SARS-CoV-2 with a 21 amino-acid C-terminal deletion^29^. The spike-expressing VSV virus was propagated in MA104 cells (African green monkey, ATCC CRL-2378.1) as described previously^29^, and viral stocks were titrated on Vero E6 cell monolayer cultures. Plaques were visualized using neutral red staining. To screen for escape mutations selected in the presence of COV2-2196, COV2-2130, or a cocktail composed of a 1:1 mixture of COV2-2196 and COV2-2130, we used a real-time cell analysis assay (RTCA) and xCELLigence RTCA MP Analyzer (ACEA Biosciences Inc.) and a previously described escape selection scheme^27^. Briefly, 50 μL of cell culture medium (DMEM supplemented with 2% FBS) was added to each well of a 96-well E-plate to obtain a background reading. Eighteen thousand (18,000) Vero E6 cells in 50 μL of cell culture medium were seeded per well, and plates were placed on the analyzer. Measurements were taken automatically every 15 min and the sensograms were visualized using RTCA software version 2.1.0 (ACEA Biosciences Inc). VSV-SARS-CoV-2 virus (5,000 plaque forming units [PFU] per well, ∼0.3 MOI) was mixed with a saturating neutralizing concentration of COV2-2196, COV2-2130, or a 1:1 mixture of COV2-2196 and COV2-2130 antibody (5 μg/mL total concentration of antibodies) in a total volume of 100 μL and incubated for 1 h at 37°C. At 16-20 h after seeding the cells, the virus-antibody mixtures were added to cell monolayers. Wells containing only virus in the absence of antibody and wells containing only Vero E6 cells in medium were included on each plate as controls. Plates were measured continuously (every 15 min) for 72 h. Escape mutations were identified by monitoring the cell index for a drop in cellular viability. To verify escape from antibody selection, wells where cytopathic effect was observed in the presence of COV2-2130 were assessed in a subsequent RTCA experiment in the presence of 10 μg/mL of COV2-2130 or COV2-2196. After confirmation of resistance of selected viruses to neutralization by COV2-2130, viral isolates were expanded on Vero E6 cells in the presence of 10 μg/mL of COV2-2130. Viral RNA was isolated using a QiAmp Viral RNA extraction kit (QIAGEN) according to manufacturer protocol, and the SARS-CoV-2 spike gene was reverse-transcribed and amplified with a SuperScript IV One-Step RT-PCR kit (ThermoFisher Scientific) using primers flanking the S gene. The amplified PCR product was purified using SPRI magnetic beads (Beckman Coulter) at a 1:1 ratio and sequenced by the Sanger method, using primers giving forward and reverse reads of the RBD.

### Serial passaging and testing of SARS-CoV-2 to select for mAb resistant mutations

SARS-CoV-2 strain USA-WA1/2020 was passaged serially in Vero cell monolayer cultures with AZD8895, AZD1061 or AZD7442, at concentrations beginning at their respective IC_50_ values and increased step-wise to their IC_90_ value with each passage. As a control, virus was passaged in the absence of antibody. Following the final passage, viruses were evaluated for susceptibility against the reciprocal antibody at a final concentration of 10 times the IC_90_ concentration by plaque assay. Plaques (n=6) were selected randomly for AZD1061 cultures, and their virus spike-encoding gene was sequenced.

### Generation of authentic SARS-CoV-2 viruses, including viruses with variant residues

The 2019n-CoV/USA_WA1/2020 isolate of SARS-CoV-2 was obtained from the US Centers for Disease Control (CDC) and passaged on Vero E6 cells. Individual point mutations in the spike gene (D614G and E484K/D614G) were introduced into an infectious cDNA clone of the 2019n-CoV/USA_WA1/2020 strain as described previously^70^. Nucleotide substitutions were introduced into a subclone puc57-CoV-2-F6 containing the spike gene of the SARS-CoV-2 wild-type infectious clone^71^. The full-length infectious cDNA clones of the variant SARS-CoV-2 viruses were assembled by *in vitro* ligation of seven contiguous cDNA fragments following the previously described protocol^71^. *In vitro* transcription then was performed to synthesize full-length genomic RNA. To recover the mutant viruses, the RNA transcripts were electroporated into Vero E6 cells. The viruses from the supernatant of cells were collected 40 h later and served as p0 stocks. All virus stocks were confirmed by sequencing.

### Focus reduction neutralization test

Serial dilutions of mAbs or serum were incubated with 10^2^ focus-forming units (FFU) of different strains or variants of SARS-CoV-2 for 1 h at 37°C. Antibody-virus complexes were added to Vero-hACE2-TMPRSS2 cell monolayer cultures in 96-well plates and incubated at 37°C for 1 h. Subsequently, cells were overlaid with 1% (w/v) methylcellulose in MEM supplemented with 2% FBS. Plates were harvested 20 h later by removing overlays and fixed with 4% PFA in PBS for 20 min at room temperature. Plates were washed and sequentially incubated with an oligoclonal pool of anti-S mAbs and HRP-conjugated goat anti-human IgG in PBS supplemented with 0.1% saponin and 0.1% bovine serum albumin. SARS-CoV-2-infected cell foci were visualized using TrueBlue peroxidase substrate (KPL) and quantitated on an ImmunoSpot microanalyzer (Cellular Technologies).

## Multiple sequence alignments

We searched for antibody variable gene sequences originating with the same features as those encoding COV2-2196 and retrieved the matching sequences from the repertoires of each individual examined. We searched for similar sequences in the publicly available large-scale antibody sequence repertoires for three healthy individuals and cord blood repertoires (deposited at SRP174305). The search parameters for the heavy chain were sequences with *IGHV1-58* and *IGHJ3* with the P99, D108, and F110 residues. Additionally, the search parameters for the light chain were sequences with Y92 and W98 residues. Sequences from a matching clonotype that belonged to each individual were aligned with either ClustalO^72^ (heavy chains) or with MUSCLE^73^ (light chains). Then, LOGOs plots of aligned sequences were generated using WebLogo^74^.

