## Extended Data Table 1 for "Genetic and structural basis for recognition of SARS-CoV-2 spike protein by a two-antibody cocktail"

| Data collection |  |  |
| --- | --- | --- |
| Crystal | RBD-COV2-2196 | RBD-COV2-2196-2130 |
| PDB ID | 7L7D | 7L7E |
| Wave Length (Å) | 0.97872 | 0.97857 |
| Space group | P2_1_ | P2_1_ |
| Unit cell dimensions |  |  |
| a, b, c (Å) | 44.1, 81.6, 101.2 | 97.2, 152.5, 199.2 |
| α, β, γ | 90.0, 96.6, 90.0 | 90.0, 94.7, 90.0 |
| Resolution (Å) | 43.98 – 2.50 | 35.21 – 3.00 |
| Unique reflections | 24896 (13417) | 115751 (16908) |
| Redundancy | 3.7 (3.7) | 7.7 (7.7) |
| Completeness (%) | 99.9 (99.6) | 99.9 (100) |
| R_merge_ (%) | 4.6 (24.5) | 23.1 (81.0) |
| I/σ(I) | 19.2 (4.8) | 6.4 (2.3) |
| Refinement statistics |  |  |
| R_factor_ (%) | 18.5 | 21.5 |
| R_free_ (%) | 23.1 | 27.3 |
| R.m.s.d. (bond) (Å) | 0.0030 | 0.0023 |
| R.m.s.d. (angle) (deg) | 0.563 | 0.598 |
| Ramachandran plot |  |  |
| Favored (%) | 95.82 | 95.34 |
| Allowed (%) | 4.18 | 4.37 |
| Outliers (%) | 0.00 | 0.09 |

**Extended Data Table 1. Data collection and refinement statistics for the crystals of RBD-COV2-2196 and RBD-COV2-2196-2130 complexes**

R_merge_ = Σ Σ |I_hkl_ − I_hkl(j)_|/Σ I_hkl_, where I_hkl(j)_ is the observed intensity and I_hkl_ is the final average intensity.

R_work_ = Σ ||Fobs| − |Fcalc|/Σ |Fobs| and R_free_ = Σ ||Fobs| − |Fcalc||/Σ |Fobs|, where R_free_ and R_work_ are calculated using a randomly selected test set of 5% of the data and all reflections excluding the 5% test set, respectively. Numbers in parentheses are for the highest resolution shell.
