## Extended Data Figures 1 to 8 for "Genetic and structural basis for recognition of SARS-CoV-2 spike protein by a two-antibody cocktail"

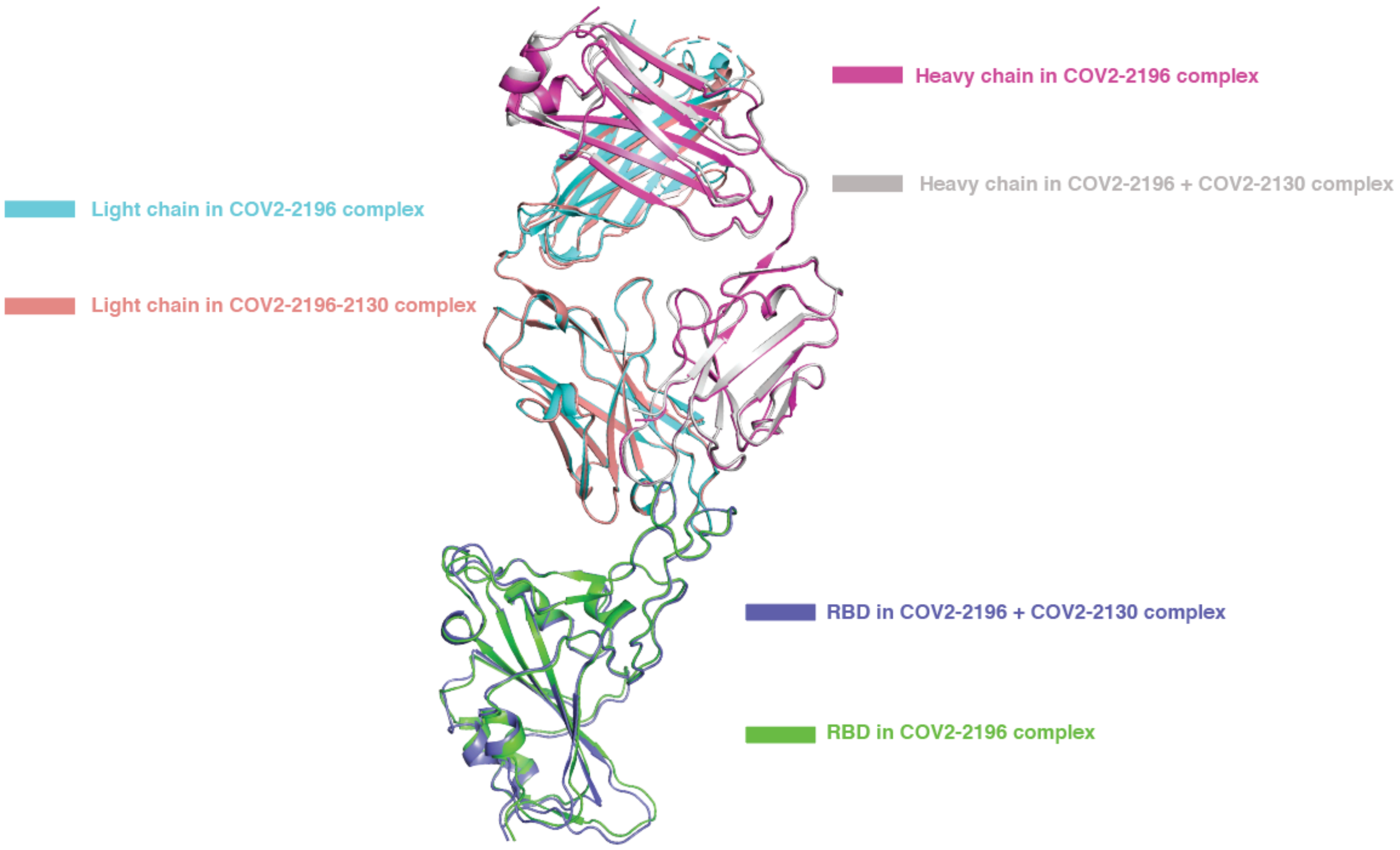


**Extended Data Fig. 1. Overlay of substructure of RBD-COV2-2196 in the RBD-COV2-2196 + COV2-2130 complex and the RBD-COV2-2196 crystal structure.**

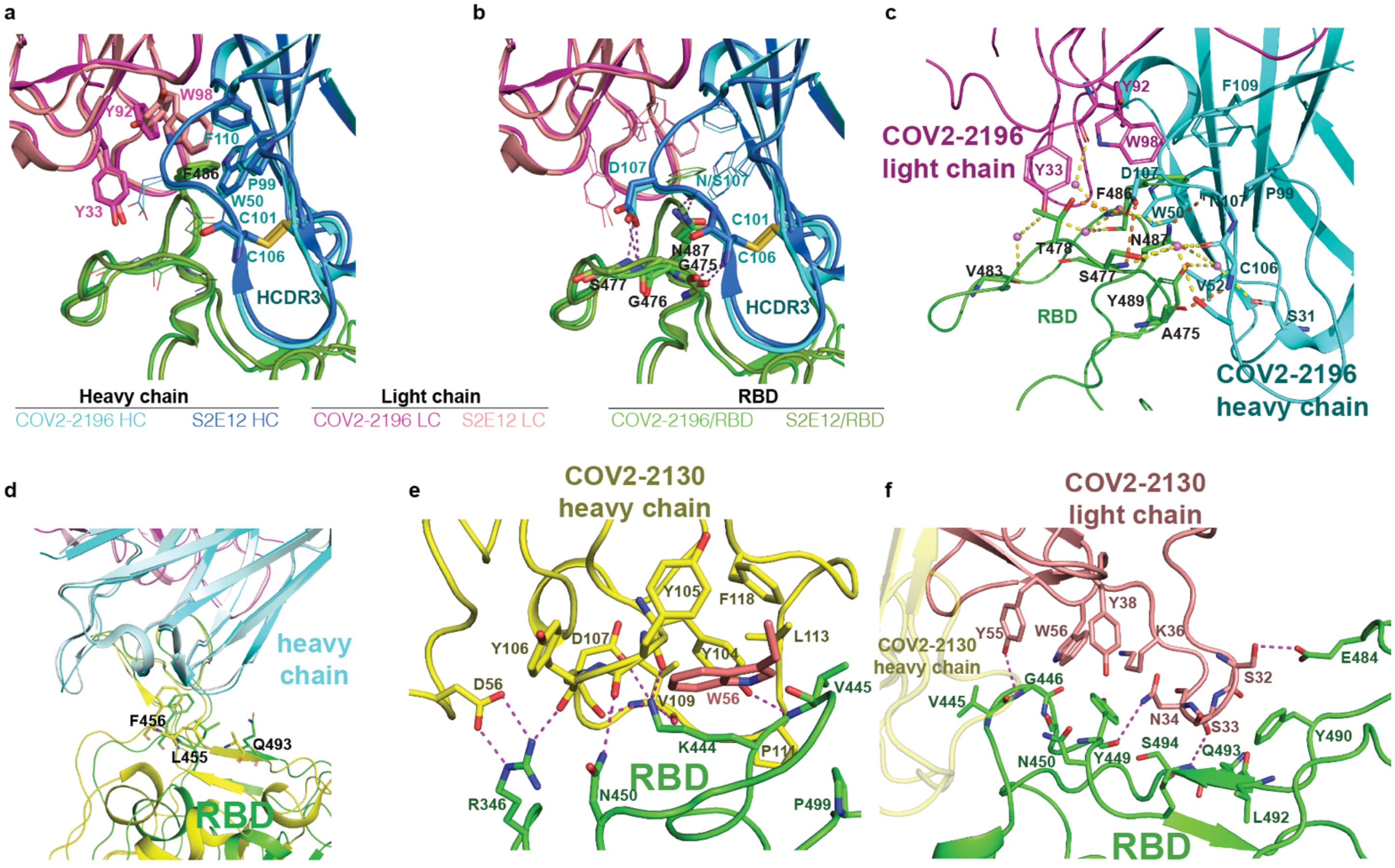


**Extended Data Fig. 2.** Similar aromatic stacking and hydrophobic interaction patterns at the RBD site F486 shared between RBD-COV2-2196 and spike-S2E12 complexes.

- **a.** Same hydrogen bonding pattern surrounding residue F486 in the structures of the two complexes.
- **b.** Detailed interactions between COV2-2196 and RBD. COV2-2196 heavy chain is colored in cyan, the light chain is colored in magenta, and RBD is colored in green. Important interacting residues are shown in stick representation. Water molecules involved in antibody-RBD interaction are represented as pink spheres. Direct hydrogen bonds are shown as orange dashed lines, and water-mediated hydrogen bonds as yellow dashed lines.
- **c.** Superimposition of S2E12/RBD cryo-EM structure onto the COV2-2196/RBD crystal structure, with the variable domains of antibodies as references. COV2-2196 heavy chain is in cyan, and its light chain in magenta; S2E12 heavy chain is in pale cyan, and its light chain in light pink. The two corresponding RBD structures are colored in green or yellow, respectively.
- **d.** Detailed interactions between COV2-2130 heavy chain and RBD. Paratope residues are shown in stick representation and colored in yellow, epitope residues in green sticks. Hydrogen-bonds or strong polar interactions are represented as dashed magenta lines.
- **e.** Detailed interactions between COV2-2130 light chain and RBD. Paratope residues are shown in stick representation and colored in orange, epitope residues in green sticks. Hydrogen-bonds are represented as dashed magenta lines.

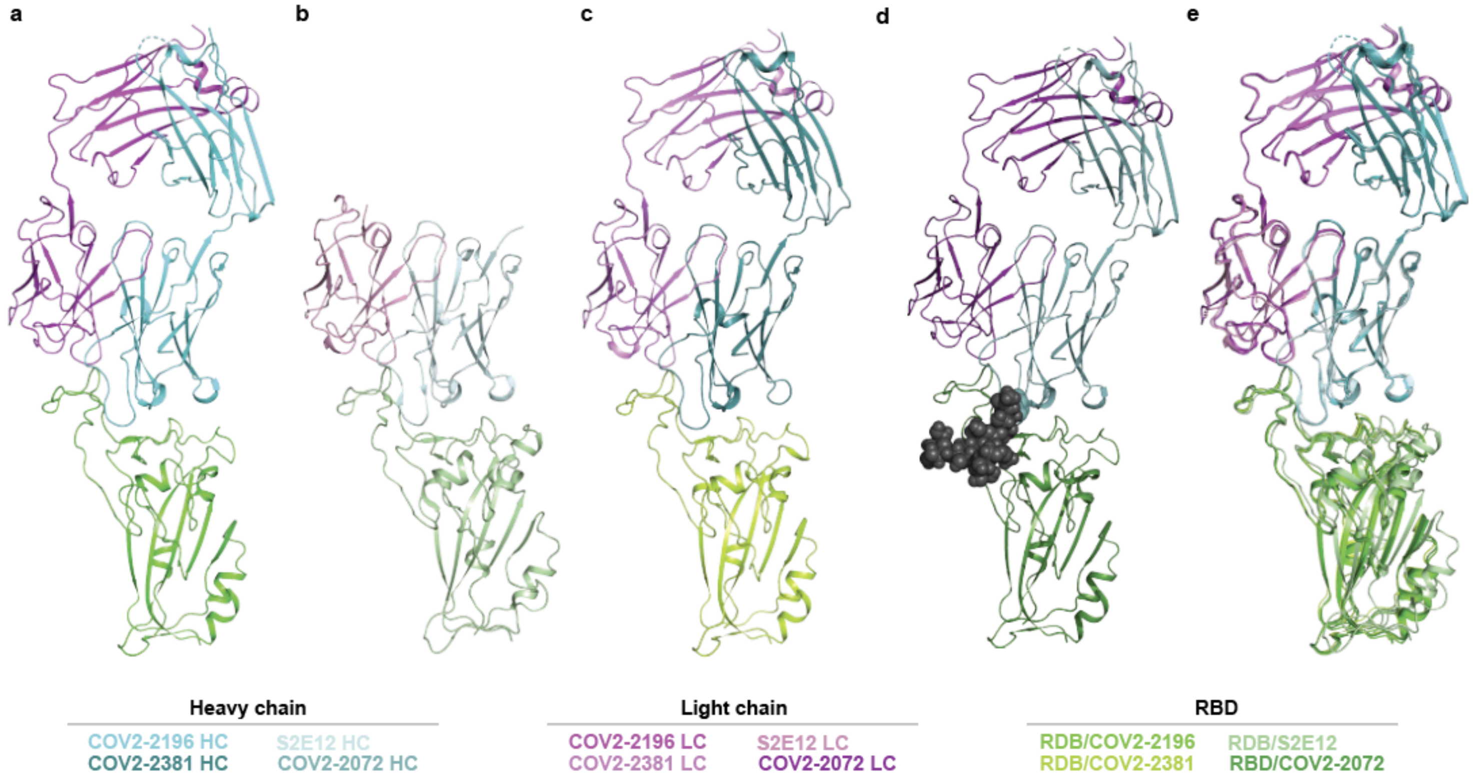


**Extended Data Fig. 3. A common clonotype of anti-RBD antibodies with the same binding mechanism.**

**a.** COV2-2196/RBD crystal structure.

**b.** S2E12/RBD cryo-EM structure.

**c.** COV2-2381/RBD homology model. COV2-2072 encodes an N-linked glycosylation sequon in the HCDR3, indicated by the gray spheres.

**d.** COV2-2072/RBD homology model.

**e.** Overlay of the COV2-2196/RBD crystal structure (**a**) and S2E12/RBD cryo-EM structure (**b**).

**a**

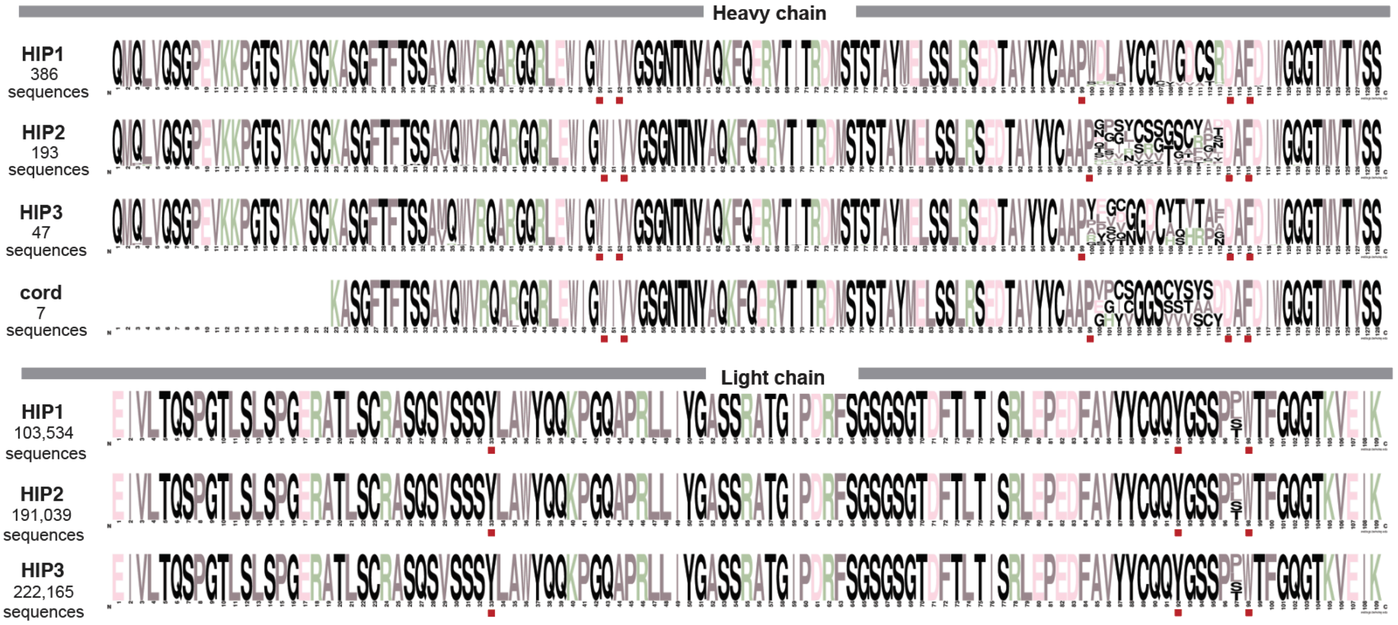

**b**

| Subject | Light chain amino acid multiple sequence alignment | Number of occurrences |
| --- | --- | --- |
| HIP1 | EIVLTQSPGTLSLSPGERATLSCRASQSVSSSYLAWYQQKPGQAPRLLIYGASSRATGIPDRFSGSGSGTDFTLTISRLEPEDFAVYYCQQYGSS-PWTFGQGTKVEIK | 2,323 |
| | EIVLTQSPGTLSLSPGERATLSCRASQSVSSSYLAWYQQKPGQAPRLLIYGASSRATGIPDRFSGSGSGTDFTLTISRLEPEDFAVYYCQQYGSSPPWTFGQGTKVEIK | 1,087 |
| | EIVLTQSPGTLSLSPGERATLSCRASQSVSSSYLAWYQQKPGQAPRLLIYGASSRATGIPDRFSGSGSGTDFTLTISRLEPEDFAVYYCQQYGSS-LWTFGQGTKVEIK | 1,086 |
| | EIVLTQSPGTLSLSPGERATLSCRASQSVSSSYLAWYQQKPGQAPRLLIYGASSRATGIPDRFSGSGSGTDFTLTISRLEPEDFAVYYCQQYGSS-SWTFGQGTKVEIK | 601 |
| | EIVLTQSPGTLSLSPGERATLSCRASQSVSSSYLAWYQQKPGQAPRLLIYGASSRATGIPDRFSGSGSGTDFTLTISRLEPEDFAVYYCQQYGSSPTWTFGQGTKVEIK | 539 |
| HIP2 | EIVLTQSPGTLSLSPGERATLSCRASQSVSSSYLAWYQQKPGQAPRLLIYGASSRATGIPDRFSGSGSGTDFTLTISRLEPEDFAVYYCQQYGSS-PWTFGQGTKVEIK | 3,909 |
| | EIVLTQSPGTLSLSPGERATLSCRASQSVSSSYLAWYQQKPGQAPRLLIYGASSRATGIPDRFSGSGSGTDFTLTISRLEPEDFAVYYCQQYGSS-LWTFGQGTKVEIK | 2,205 |
| | EIVLTQSPGTLSLSPGERATLSCRASQSVSSSYLAWYQQKPGQAPRLLIYGASSRATGIPDRFSGSGSGTDFTLTISRLEPEDFAVYYCQQYGSSPPWTFGQGTKVEIK | 1,747 |
| | EIVLTQSPGTLSLSPGERATLSCRASQSVSSSYLAWYQQKPGQAPRLLIYGASSRATGIPDRFSGSGSGTDFTLTISRLEPEDFAVYYCQQYGSS-SWTFGQGTKVEIK | 1,246 |
| | EIVLTQSPGTLSLSPGERATLSCRASQSVSSSYLAWYQQKPGQAPRLLIYGASSRATGIPDRFSGSGSGTDFTLTISRLEPEDFAVYYCQQYGSSPTWTFGQGTKVEIK | 1,230 |
| HIP3 | EIVLTQSPGTLSLSPGERATLSCRASQSVSSSYLAWYQQKPGQAPRLLIYGASSRATGIPDRFSGSGSGTDFTLTISRLEPEDFAVYYCQQYGSS-PWTFGQGTKVEIK | 3,908 |
| | EIVLTQSPGTLSLSPGERATLSCRASQSVSSSYLAWYQQKPGQAPRLLIYGASSRATGIPDRFSGSGSGTDFTLTISRLEPEDFAVYYCQQYGSS-LWTFGQGTKVEIK | 1,889 |
| | EIVLTQSPGTLSLSPGERATLSCRASQSVSSSYLAWYQQKPGQAPRLLIYGASSRATGIPDRFSGSGSGTDFTLTISRLEPEDFAVYYCQQYGSSPPWTFGQGTKVEIK | 1,809 |
| | EIVLTQSPGTLSLSPGERATLSCRASQSVSSSYLAWYQQKPGQAPRLLIYGASSRATGIPDRFSGSGSGTDFTLTISRLEPEDFAVYYCQQYGSS-SWTFGQGTKVEIK | 1,293 |
| | EIVLTQSPGTLSLSPGERATLSCRASQSVSSSYLAWYQQKPGQAPRLLIYGASSRATGIPDRFSGSGSGTDFTLTISRLEPEDFAVYYCQQYGSSPTWTFGQGTKVEIK | 953 |

**Extended Data Fig. 4. Identification of putative public clonotype members genetically similar to COV2-2196 in the antibody variable gene repertoires of virus-naïve individuals.** Antibody variable gene sequences collected from healthy individuals prior to the pandemic with the same sequence features as COV2-2196 heavy chain (**a**) and light chain (**b**) are aligned. Sequences from three different donors as well as cord blood included sequences with the features of the public clonotype. The sequence features and contact residues used in COV2-2196 are highlighted in red boxes below each multiple sequence alignment.

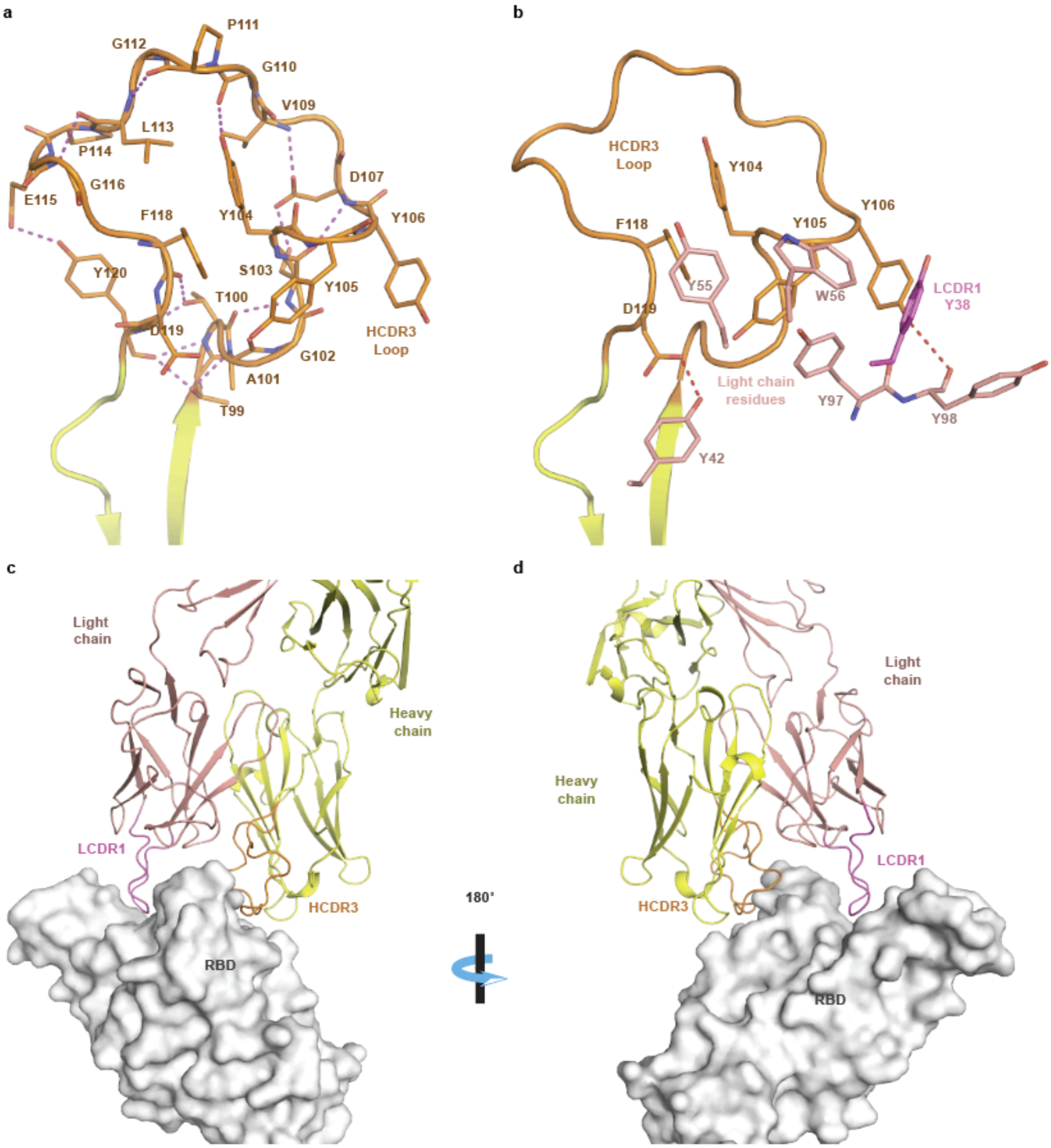


**Extended Data Fig. 5.**

- **a.** Detailed COV2-2130 HCDR3 loop structure. Short-range hydrogen bonds, stabilizing the loop conformation, are shown as dashed magenta lines.
- **b.** Residues of COV2-2130 light chain form aromatic stacking interactions and hydrogen bonds with HCDR3 to further stabilize the HCDR3 loop.
- **c.** Long LCDR1, HCDR2, and HCDR3 form complementary binding surface to the RBD epitope. RBD is shown as surface representation in grey. COV2-2130 heavy chain is colored in yellow with HCDR3 in orange, and the light chain in salmon with LCDR1 in magenta.
- **d.** 180° rotation view of panel **c**.

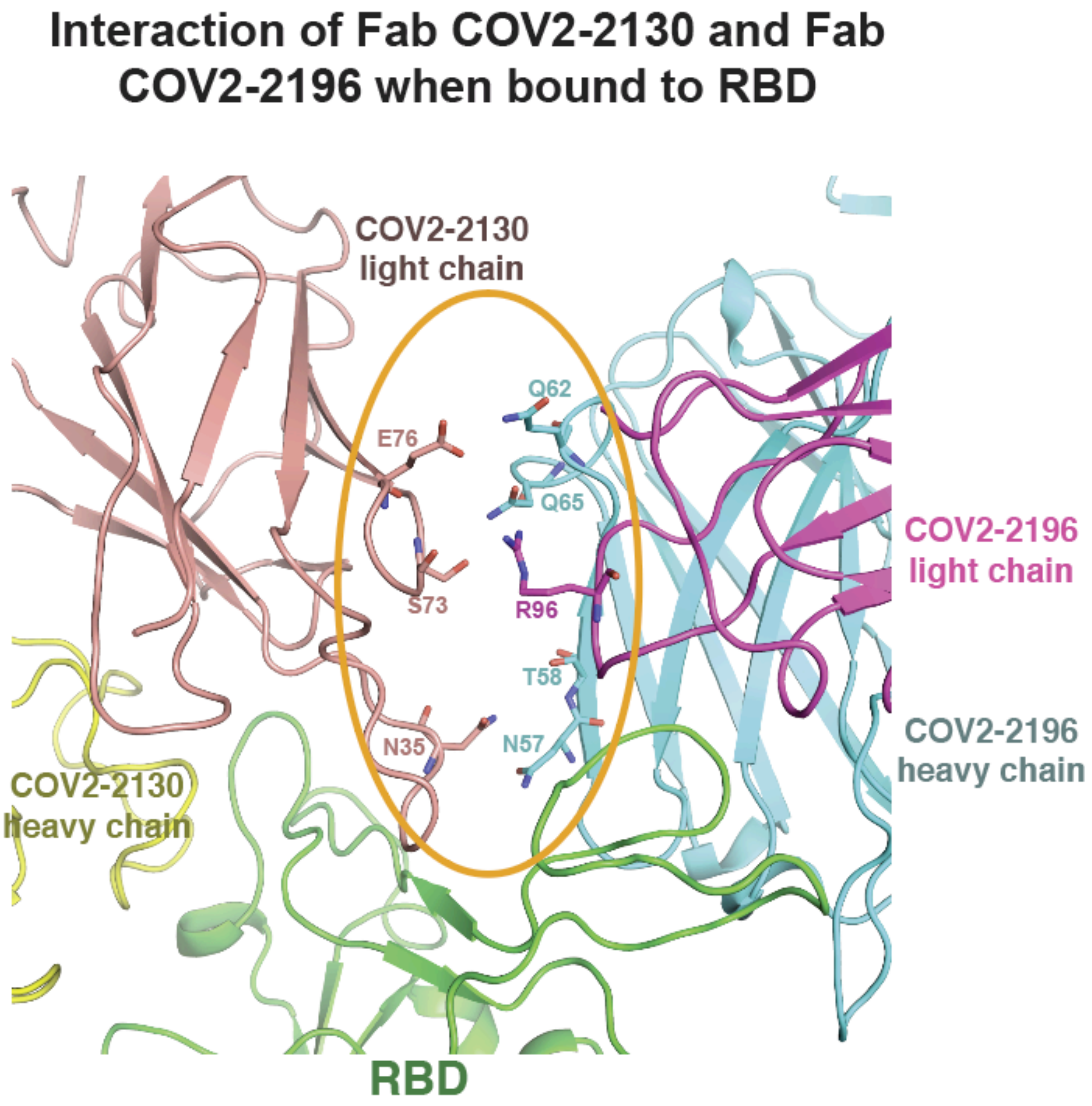


**Extended Data Fig. 6. Interface between COV2-2196 and COV2-2130 in the crystal structure of RBD in complex with COV2-2196 and COV2-2130.** COV2-2196 heavy or light chain are shown as cartoon representation in cyan or magenta, respectively, and COV2-2130 heavy or light chain in yellow or salmon, respectively. The RBD is colored in green. Interface residues are shown in stick representation.

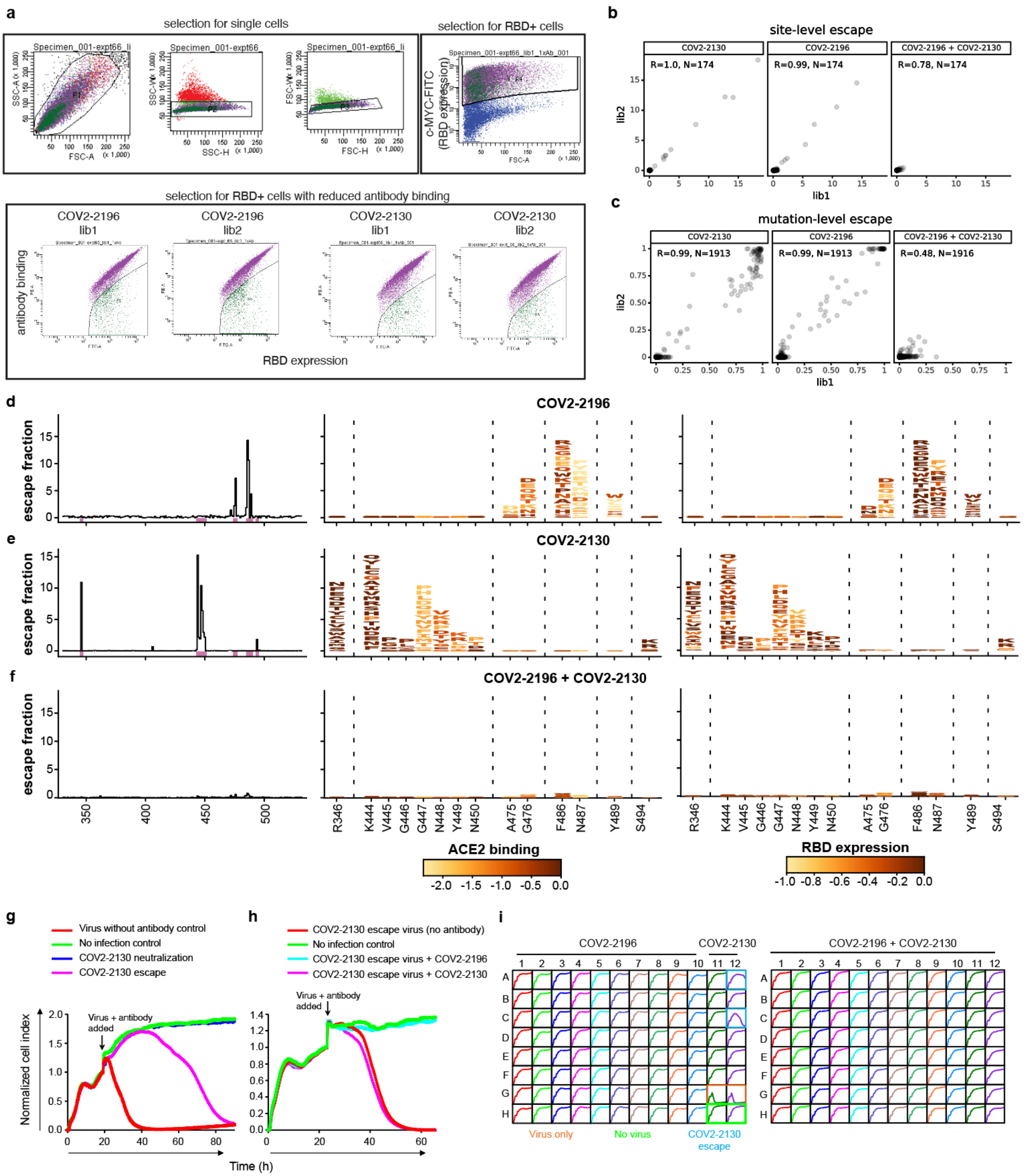


**Extended Data Fig. 7. Identification by deep mutational scanning of mutations affecting antibody binding and method of selection of antibody resistant mutants with VSV-SARS-CoV-2 virus.**

**a.** Top: Flow cytometry plots showing representative gating strategy for selection of single yeast cells using forward- and side-scatter (first three panels) and selection of yeast cells expressing RBD (right panel). Each plot is derived from the preceding gate. Bottom: Flow cytometry plots showing gating for RBD$^{+}$, antibody⁻ yeast cells (*i.e.*, cells that express RBD but where a mutation prevents antibody binding). Selection experiments are shown for COV2-2196 or COV2-2130, with two independent libraries shown for each.

**b.** Correlation of observed sites of escape from antibody binding between yeast library selection experiments using COV2-2196, COV2-2130, or a 1:1 mixture of COV2-2196 and COV2-2130. The x-axes show cumulative escape fraction for each site for library 1, and the y-axes show cumulative escape fraction for each site for library 2. Correlation coefficient and *n* are denoted for each graph.

**c.** Correlation of observed mutations that escape antibody binding between yeast library selection experiments using COV2-2196, COV2-2130, or a 1:1 mixture of COV2-2196 and COV2-2130. The x-axes show each amino acid mutation's escape fraction for library 1, and the y-axes show each amino acid mutation's escape fraction for library 2. Correlation coefficient and *n* are denoted for each graph.

**d-f.** DMS results for COV2-2196 (**d**), COV2-2130 (**e**), or a 1:1 mixture of COV2-2196 and COV2 2130 (**f**). Left panels: sites of escape across the entire RBD are indicated by peaks that correspond to the logo plots in the middle and right panel. Middle panel: as in **Fig. 4a**, logo plot of cumulative escape mutation fractions of all RBD sites with strong escape mutations for COV2-2196, or COV2-2130, or COV2-2196+COV2-2130. Mutations are colored based on the degree to which they abrogate RBD binding to human ACE2. Right panel: again, logo plots show cumulative escape fractions, but colored based on the degree to which mutations effect RBD expression in the yeast display system. Interactive, zoomable versions of these logo plots are at https://jbloomlab.github.io/SARS-CoV-2-RBD_MAP_AZ_Abs/.

**g.** Representative RTCA sensograms showing virus that escaped antibody neutralization. Cytopathic effect (CPE) was monitored kinetically in Vero E6 cells inoculated with virus in the presence of a saturating concentration (5 μg/mL) of antibody COV2-2130. Representative instances of escape (magenta) or lack of detectable escape (blue) are shown. Uninfected cells (green) or cells inoculated with virus without antibody (red) serve as controls. Magenta and blue curves represent a single representative well; the red and green controls are the mean of technical duplicates.

**h.** Representative RTCA sensograms validating that a variant virus selected by COV2-2130 in (**g**) indeed escaped COV2-2130 (magenta) but was neutralized by COV2-2196 (light blue).

**i.** Example sensograms from individual wells of 96-well E-plate analysis for escape selection experiments with COV2-2196, COV2-2130, or a 1:1 mix of COV2-2196 and COV2-2130. Instances of escape from COV2-2130 are noted, while escape was not detected in the presence of COV2-2196 or COV2-2196+COV2-2130. Positive and negative controls are denoted on the first plate.

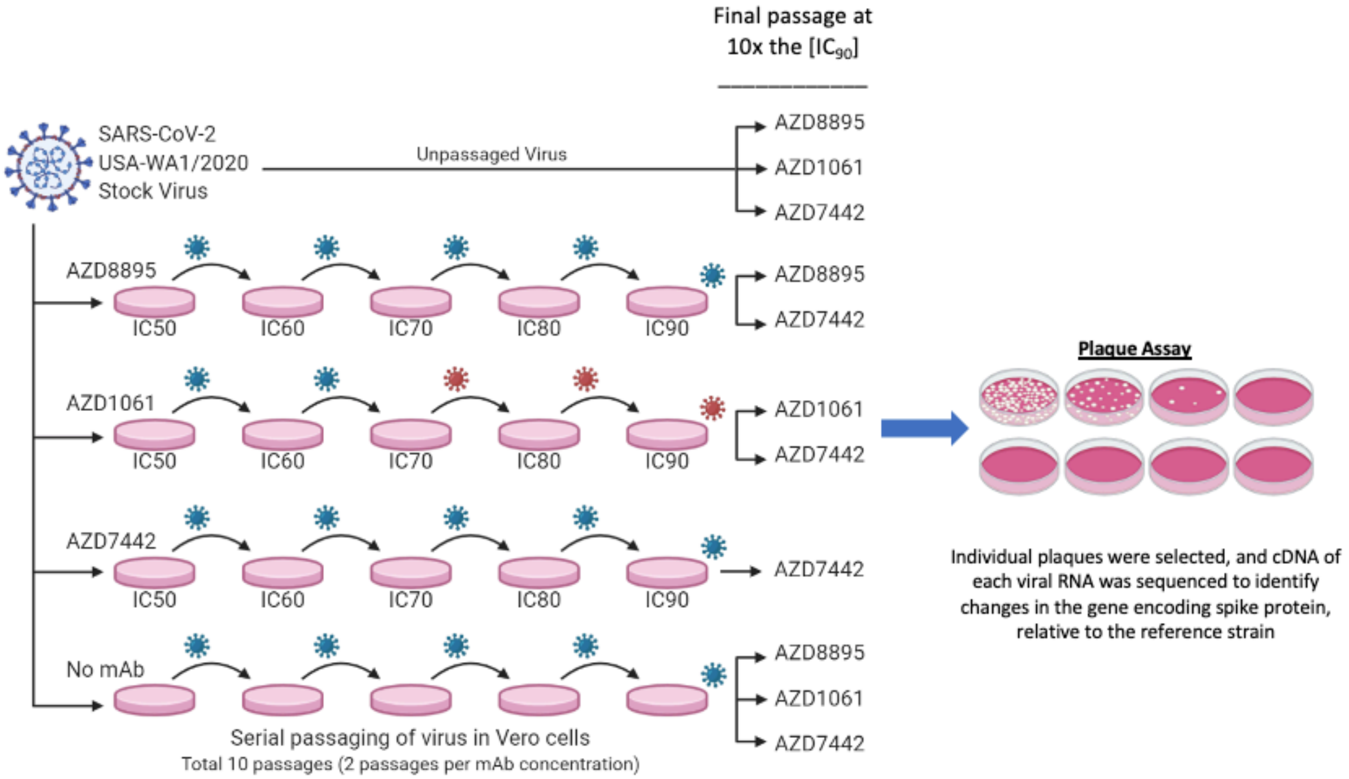


**Extended Data Fig. 8. Method of selection of antibody resistant mutants with authentic SARS-CoV-2 virus.** The method for assessing mAb-resistant spike protein variants is shown. SARS-CoV-2 was passaged serially in the presence of mAbs at the increasing concentrations indicated in the figure or without antibody (no mAb). Following passage at $IC_{90}$ concentrations, samples were treated with 10× the $IC_{90}$ concentrations of mAbs and any resultant resistant virus collected, and the genome was sequenced.
